# A robust and comprehensive quality control of cerebral cortical organoids: methodology and validation

**DOI:** 10.1101/2025.03.13.642794

**Authors:** Héloïse Castiglione, Lucie Madrange, Camille Baquerre, Benoît Guy Christian Maisonneuve, Jean-Philippe Deslys, Frank Yates, Thibault Honegger, Jessica Rontard, Pierre-Antoine Vigneron

## Abstract

Cerebral organoids hold great promise for neuroscience research as complex *in vitro* models that mimic human brain development. However, they face significant challenges related to quality and reproducibility, leading to unreliability in both academic and industrial contexts. Discrepancies in morphology, size, cellular composition, and cytoarchitectural organization limit their application in biomedical studies, particularly in disease modeling, drug screening, and neurotoxicity testing, where consistent models are essential. Critically, current methods for organoid characterization often lack standardization and rely heavily on subjective assessments, restricting their broader applicability. In this study, we developed a comprehensive Quality Control (QC) framework for 60-days cortical organoids. Five key criteria: morphology, size and growth profile, cellular composition, cytoarchitectural organization, and cytotoxicity, are evaluated using a standardized scoring system. We implemented a hierarchical approach, beginning with non-invasive assessments to exclude low-quality organoids (Initial Scoring), while reserving in-depth analyses for those that passed the initial evaluation (Final Scoring). To validate this framework, we exposed 60-day cortical organoids to graded doses of hydrogen peroxide (H_2_O_2_), inducing a spectrum of quality outcomes. The QC system demonstrated its robustness and reproducibility by accurately discriminating organoid quality based on objective and quantifiable metrics. This standardized and user-friendly framework for quality assessment not only minimizes observer bias but also enhances the reliability and comparability of cerebral organoid studies. Additionally, its scalability makes it suitable for industrial applications and adaptable to other organoid types, offering a valuable tool for advancing both fundamental and preclinical research.

## INTRODUCTION

Cerebral organoids have emerged as innovative tools in neuroscience by providing biologically relevant *in vitro* models that recapitulate aspects of the human brain development and function. These three-dimensional (3D) structures, derived from the neuroectodermal differentiation of pluripotent stem cells, self-organize into complex architectures recapitulating certain regions of the human brain [1], such as the forebrain, midbrain, hindbrain, or even more specifically the hippocampus, cortex, or choroid plexus [2–12]. Unspecific differentiation protocol can also give rise to unguided whole-brain organoids [13].

Unlike traditional 2D cultures or simpler 3D models such as spheroids and neurospheres, cerebral organoids recreate a physiologically relevant cellular microenvironment. This complexity allows for enhanced cell-cell and cell-matrix interactions, fostering improved differentiation and maturation [14]. While human brain organogenesis remains a highly complex process, tightly regulated both on a spatial and a temporal scale [15], cerebral organoids have proven their ability to model key neurodevelopmental aspects, including neurogenesis, neuronal migration, neuromorphogenesis and synaptogenesis [1, 15]. Furthermore, transcriptomic and epigenetic analyses have revealed that these models closely mimic developmental trajectories observed in the human fetal brain [15, 16]. When derived from patient-specific cells, or when combined with advanced genetic engineering techniques, such features have made cerebral organoids powerful tools for studying neurodevelopmental disorders, such as microcephaly [13] and trisomy 21 [17–19], as well as for studying neurological cancers [20], and can also give clues about the pathogenesis of neurodegenerative diseases, including Alzheimer’s disease [21, 22], Parkinson’s disease [23], and Creutzfeldt-Jakob disease [24].

Beyond modeling diseases, cerebral organoids have shown promise in neurotoxicity studies [25, 26]. Notably, the developing human brain is highly susceptible to environmental insults, and exposure to pollutants or chemicals during pregnancy can disrupt its physiological development. Organoids could provide an unprecedented human-based predictive model to study developmental neurotoxicity (DNT) in response to drugs, chemicals, and pollutants. Studies using cerebral organoids have already explored the effects of valproic acid [27–34], nicotine [35], cannabis [36], bisphenols [37, 38], cadmium [39] and nanoplastics [40], among others [41–44].

Despite their potential, cerebral organoids face significant challenges in terms of quality and reproducibility. Morphological inconsistencies, variations in size, and differences in cellular composition or cytoarchitectural organization often arise from the stochastic nature of stem cell differentiation and the spontaneous self-organization occurring within the organoid [1, 45]. For instance, within a batch of cerebral organoids, some organoids will display optimal morphology, with dense overall structure and well-defined borders, while others maybe poorly compact and will tend to degrade over time by losing cells [46, 47]. Moreover, some organoids will exhibit expected cell types and cytoarchitectural organization, whereas others may present disorganized structures and lower proportions of some cell types. Similarly, suboptimal cystic cavities can also be present within some organoids or protrude from their surface [46]. In addition, a necrotic core can also arise in certain organoids [1, 45, 48]. Non-cerebral structures might also occasionally occur, including germ layers other than neuroectoderm, especially in the unguided-differentiated organoids [1, 49]. These inconsistencies compromise the reproducibility of scientific results, particularly in disease modeling, neurotoxicity testing, and preclinical drug screening, where high-quality and consistent models are essential [45]. Furthermore, the lack of standardized criteria for organoid generation, culture, and characterization exacerbates this variability, creating barriers to their broader adoption in industrial and preclinical applications.

Current methods for organoid characterization, including immunohistochemistry [2, 13], transcriptomic profiling [6], electrophysiological recording [50], and cytotoxicity studies [51–55] are valuable but often lack standardization and face several limitations. Many current approaches rely on qualitative and subjective assessments that might introduce inconsistencies and bias. It is common that for daily evaluation of cerebral organoids, researchers rely on morphological observations to assess quality, but this qualitative readout is not frequently detailed in research publications. Although morphological criteria are often used and provide valuable information, their translation into standardized quantitative indicators transferable between laboratories remains partially done, even if recent publications highlight a growing interest in leveraging these criteria as reliable, non-invasive readouts for characterizing cerebral organoids [56–58]. Moreover, some analysis methods commonly used in 2D cell cultures are difficult to transpose to 3D cultures, further complicating the standardization of their characterization [55, 59]. Overall, there is a notable lack of robust and well-defined quantitative methodologies for 3D organoid characterization. This gap limits the ability to objectively evaluate cerebral organoids in terms of quality, especially across diverse research groups, ultimately affecting the reliability and consistency of results.

In this study, we propose a comprehensive and robust Quality Control (QC) framework for 60-day cortical organoids to address these challenges in their evaluation. This system integrates five critical criteria: A) Morphology, B) Size and Growth Profile, C) Cellular Composition, D) Cytoarchitectural Organization, and E) Cytotoxicity, into a standardized scoring methodology. The framework is designed hierarchically, prioritizing early, non-invasive evaluations to efficiently exclude organoids of low quality, while reserving in-depth analyses for organoids that have met initial thresholds. To validate its reliability and applicability, we exposed 60-day cortical organoids to gradual doses of hydrogen peroxide (H_2_O_2_), producing varying quality levels to rigorously test the scoring system. By minimizing observer bias and enabling objective, reproducible quality assessments, this QC framework enhances the consistency and comparability of results in cerebral organoid research. Moreover, its potential to support both academic studies and industrial scalability highlights its value as a versatile tool for advancing biomedical research.

## MATERIALS AND METHODS

### hiPSC culture and maintenance

Human induced Pluripotent Stem Cells (hiPSCs) were generated by reprogramming BJ primary foreskin fibroblasts obtained from ATCC (CRL-2522), using the non-integrative Sendai virus vectors following the manufacturer’s instructions (A16517, ThermoFisher Scientific). Pluripotency was confirmed by identifying specific pluripotency markers through Reverse Transcriptase-Polymerase Chain Reaction (RT-PCR), and regular tests were conducted to verify the absence of mycoplasma. The culture and maintenance of hiPSCs were performed as previously reported [21, 22, 55]. Briefly, hiPSCs were maintained on Geltrex-coated cell culture plates (A1569601, Gibco) and cultured in mTeSR™ Plus medium (100-0276, STEMCELL Technologies) supplemented with 1% Penicillin/Streptomycin (P/S) (15140122, Gibco), at 37°C in a 5%-enriched CO_2_ atmosphere. hiPSCs were passaged upon reaching 50-70% confluency using 0.02% ethylenediaminetetraacetic acid (EDTA) treatment (E8008, Sigma-Aldrich).

### Generation and culture of cerebral cortical organoids

Cerebral cortical organoids were generated as previously reported [55], from a protocol adapted from methods described by Xiang *et al*. [4, 5] relying on dorsal forebrain-regionalized differentiation. On day 0, hiPSCs were detached using 0.02% EDTA treatment and dissociated with Accutase (AT-104, STEMCELL Technologies) to obtain a single-cell suspension. These cells were seeded in V-bottom cell-repellent 96-well plates (651970, Greiner Bio-One) at a density of 20,000 cells/well in neural induction medium containing Dulbecco’s Modified Eagle Medium/Nutrient Mixture F-12, GlutaMAX supplement (DMEM/F-12, 10565018, Gibco), 15% (v/v) KnockOut Serum Replacement (KOSR, 10828010, Gibco), 1% Minimum Essential Medium-Non-Essential Amino Acids (MEM-NEAA, 1140035, Gibco), 1% P/S, 100 nM LDN-193189 (72147, STEMCELL Technologies), 10 μM SB-431542 (72232, STEMCELL Technologies), 2 μM XAV-939 (X3004, Sigma-Aldrich), 100 μM β-mercaptoethanol (21985023, Gibco), and supplemented with 5% Fetal Bovine Serum (FBS, 10270106, Gibco) and 50 μM Y-27632 (72304, STEMCELL Technologies). On day 2, embryoid bodies (EBs) were collected and transferred into 24-well suspension cell culture plates (144530, Nunc). The neural induction medium was renewed every two days until day 10, with FBS removed from day 2, and Y-27632 removed from day 4. From day 10 to day 18, EBs were cultured in differentiation medium without vitamin A, containing DMEM/F-12:NeuroBasal Medium (21103049, Gibco) at 1:1 ratio, supplemented with 0.5% (v/v) MEM-NEAA, 1% P/S, 0.5% N2 supplement 100X (17502-048, Gibco), 1% B-27 supplement minus vitamin A (12587010, Gibco), 1% HEPES solution (H0887, Sigma-Aldrich), 0.025% human insulin (19278-5 mL, Sigma-Aldrich), and 50 μM β-mercaptoethanol. From day 18, EBs were cultured in a differentiation medium with vitamin A, following the same composition as the previously described medium, but replacing the B-27 supplement minus vitamin A, with B-27 supplement with vitamin A (17504044, Gibco), and supplemented with 20 ng/mL BDNF (78005, STEMCELL Technologies), 200 μM ascorbic acid (A9290225G, Sigma-Aldrich), and 200 μM cAMP (73886, STEMCELL Technologies). Cortical organoids were cultured in 24-well plates with 500 μL of culture medium renewed every two days from day 2 to day 28. After day 28, the medium volume was increased to 1 mL and renewed once a week. Organoids were cultured under agitation (80 rpm/min) at 37°C, in a 5%-enriched CO_2_ atmosphere.

### H_2_O_2_ exposures on cortical organoids

Cortical organoids were exposed on day 61 of culture to hydrogen peroxide (H_2_O_2_) (1.07209.0250, Supelco) diluted in differentiation medium with vitamin A, at several doses (0.1%, 0.25%, 0.5%, 1% and 5%) during 30 min at 37°C. After exposure, cortical organoids were washed once with fresh culture medium to remove excess H_2_O_2_, were maintained in culture for 7 days, and were fixed on day 68.

### Quality Control of cortical organoids: scoring system

A multi-criteria scoring system was developed for the QC of 60-day cortical organoids (Fig. 1 and Fig. S1). It is designed with criteria following a strict hierarchy, where the most critical ones are assessed first. If an organoid fails to meet the required score for an index, it is immediately excluded from further consideration (Fig. 1A). For the QC scoring, a detailed description (Fig. S1) outlines expected values and scoring thresholds for each index. Additionally, a summary table with the minimum scores required to pass the QC for each criterion is presented (Fig. 1B). Based on their individual QC scores obtained, cortical organoids are classified into “QC passed’ or “QC failed” categories, with the specification of the failed scoring step for any organoid that did not pass the QC. More precisely, the scoring approach is tailored to be usable both for pre- and post-study, referred to as Initial QC and Final QC, respectively.

**Fig. 1:**
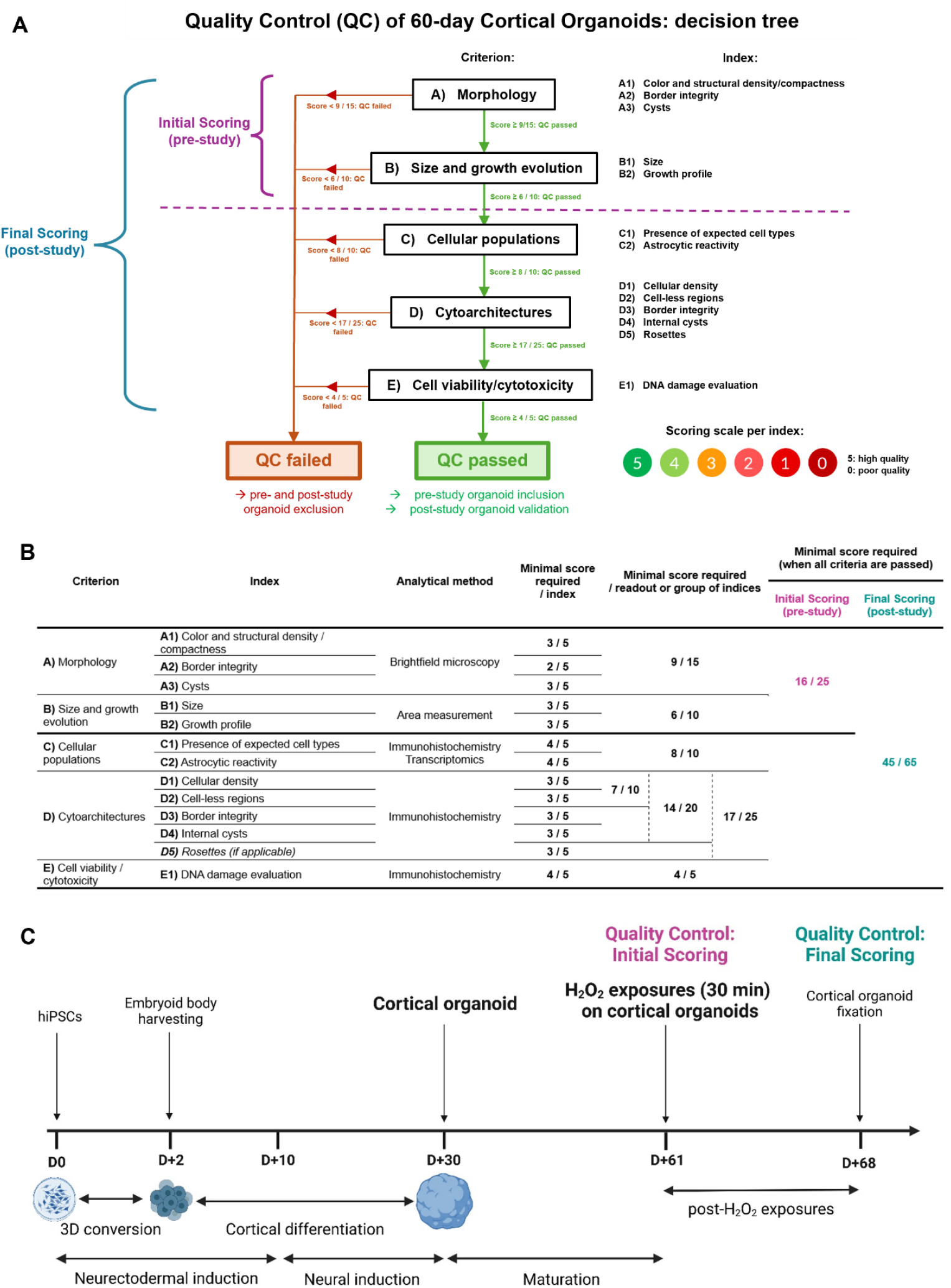
Overview of Quality Control methodology for 60-day cortical organoids, and validation by scoring of H_2_O_2_-exposed cortical organoids. (A) Overview of the Quality Control (QC) adapted to 60-day cortical organoids. The QC relies on several criteria, subdivided into indices, for cortical organoid analysis, including Morphology, Size and growth profile, Presence of expected cellular populations at 60 days, Cytoarchitectural organization, and Cellular viability and cytotoxicity levels. For each index, a scoring system enables the evaluation of organoids based on the attribution of scores comprised between 0 (poor quality) and 5 (high quality). The QC follows a hierarchical structure: criteria are assessed sequentially, and failure to meet an initial criterion automatically classifies the organoid as low quality and subsequent criteria are not assessed. The scoring system is divided into two QC: 1) an Initial Scoring to select cortical organoids before entering a study, based on the two first non-invasive criteria Morphology and Growth; and 2) a Final Scoring for complete analysis of cortical organoids based on all the criteria at the end of a study. (B) Summary table of QC criteria and minimal scores required per indices, and per composite groups of indices and readouts, that have to be obtained for cortical organoids to pass the QC. (C) Timeline of cortical organoid generation and culture protocols, including an overview of H_2_O_2_ exposure conditions. Before H_2_O_2_ exposure at D+61, cortical organoids were selected based on the Initial Scoring for QC. Exposed cortical organoids at D+68 were then evaluated following the Final Scoring for complete QC

### Initial Quality Control

In the pre-study phase, the first two non-invasive criteria, A (Morphology) and B (Size and Growth Profile), are evaluated across a batch of cortical organoids, to identify those suitable for inclusion in the subsequent study. The morphology of the organoids is assessed based on their color, density, compactness, border integrity, as well as depending on the absence or presence of cysts. In addition, organoid sizes and growth profiles are monitored to ensure they remain within expected growth ranges.

### Final Quality Control

In the post-study phase, all five criteria – A to E – are used to thoroughly validate organoid quality. This includes additional evaluations of cellular populations (criterion C), where the presence of the three expected cell types (neurons, astrocytes, neural progenitors) and astrocytic reactivity are analyzed, as well as assessments of the cytoarchitectural organization (criterion D), including cell density, proportion of cell-less regions, border integrity, presence of neurogenic areas, and occurrence of internal cysts. Finally, cell viability and cytotoxic markers (criterion E) are evaluated, with a focus on DNA damage and apoptotic markers, to ensure organoids have maintained low cytotoxicity levels throughout the study.

### Longitudinal monitoring of cortical organoid morphology and growth evolution

For cortical organoid morphology and growth profile monitoring over time, brightfield images of the organoids were acquired at regular timepoints during the culture (D+2, D+9, D+16, D+23, D+30, D+33, D+40, D+48, D+54 and D+61), using a DM IL LED Inverted Laboratory Microscope (Leica Microsystems) (5X). To assess the organoid size, the surface area of the organoid was measured from the brightfield images on FIJI/ImageJ software, version 1.54f [60].

### Immunohistochemistry

Cortical organoids were fixed in 4% paraformaldehyde (11699408, Q Path) for 2 h at room temperature (RT) under smooth agitation, followed by three washes of 10 min with Phosphate Buffered Saline solution (PBS) (18912-014, Gibco) at RT under smooth agitation, and immersed in 30% (v/v) sucrose (S9378, Sigma-Aldrich) dissolved in PBS for 48 h at 4°C. The organoids were then transferred in a solution composed of 7.5% (v/v) gelatin (G9391, Sigma-Aldrich) and 15% (v/v) sucrose dissolved in PBS for 1 h at 37°C, before being embedded in this solution for 15 min at 4°C. Embedded organoids were then snap-frozen in isopentane (M32631, Sigma-Aldrich) and stored at -80°C until use. Frozen organoids were sectioned in slices of 20 μm thickness using a cryostat (CM1850 UV, Leica Biosystems). For the immunofluorescent staining, organoid slices were permeabilized and blocked with a solution containing 0.2% Triton® X-100 (T-9284, Sigma-Aldrich), 3% bovine serum albumin (BSA, A2153, Sigma-Aldrich), and 1% normal goat serum (NGS, G9023, Sigma-Aldrich) in PBS for 1 h at RT. Then, the slices were incubated with primary antibodies diluted in the blocking solution at 4°C overnight in a humidified chamber and were washed with 0.2% Triton in PBS three times. Then, organoid slices were incubated with secondary antibodies and 4′,6-diamidino-2-phenylindole (DAPI, dilution 1:1000) for 1 h at RT in a dark humidified chamber and were washed three times with 0.2% Triton in PBS. The slices were mounted using ProLong Gold Antifade Mountant (11539306, Invitrogen), and observed under a Leica THUNDER microscope (THUNDER Imager Live Cell & 3D Assay, Leica Microsystems). Primary and secondary antibodies used are listed in Supplementary Table S1.

### Image-based quantifications of cellular density and cell-less regions

Cellular density was calculated based on DAPI positive surface, without considering cell-less zones (“holes”) since this second parameter was considered in the subsequent index. Both quantifications of cellular density and cell-less regions relied on determination of DAPI positive surface and were normalized to the total surface area of the organoid slice. Briefly, the DAPI positive surface was determined using the “Adjust Threshold” function of FIJI/ImageJ. For the cellular density, the threshold was adjusted to correspond with the DAPI labelling, while for the cell-less areas, the threshold was increased to cover the entire surface except the cell-less regions/holes. Estimation of cell number was calculated based on DAPI positive surface, considering an average nucleus area of 80 μm^2^.

### Image-based quantification of GFAP positive surface expression

Glial Fibrillary Acidic Protein (GFAP) positive surface expression was calculated using the “Adjust Threshold” function on FIJI/ImageJ, as previously described in this section, and normalized to the total DAPI positive surface area of the cryosection.

### Image-based quantification of γH2AX marker and comparison with a positive control

Quantification was conducted by counting γH2AX punctate and by normalizing to the DAPI positive surface expression, across several sections of individual organoids (n=2-3 slices per organoid, n=4 organoids per condition). Images were first binarized on FIJI/ImageJ, then the “Analyze Particles” function was applied with the following parameters: Size (micron2): 15-infinity, Circularity: 0.00-1.00. To statistically compare the maximal γH2AX immunolabelling in cerebral organoids with positive controls, the standard deviation of the positive controls was first calculated. Then, the difference between the mean value of the positive controls and the value for the organoids was determined. Finally, this difference was expressed as a ratio relative to the standard deviation of the positive controls.

## RESULTS

### Quality Control Enables Classification of 60-Day Cortical Organoids by Quality Level

We developed a comprehensive QC framework based on a scoring methodology adapted to 60-day cortical organoid evaluation and classification (Fig. 1). This QC scoring system is structured around five primary criteria (A to E) corresponding to cortical organoid analysis readouts – A) Morphology, B) Size and growth profile, C) Cellular composition, D) Cytoarchitectural organization, and E) Cytotoxicity level – each further subdivided into specific indices (Fig. 1). For each index, cortical organoids are evaluated on a scale of scores between 0 (low quality) and 5 (high quality). To streamline the process, the criteria are hierarchically organized, prioritizing non-invasive and critical assessments first (Fig. 1A). Thresholds with minimum scores are defined for each criterion (Fig. 1B), and failure to meet these thresholds halts further QC evaluation, categorizing the organoid as low-quality and resulting in its exclusion from the study. In cases where all minimal scores are achieved for a criterion, additional composite thresholds, integrating multiple indices, are applied to ensure a robust quality classification (Fig. 1B). For a detailed, illustrated and easy-to-use version of the QC scoring, see Fig. S1 in Supplementary Data.

This scoring system is designed for two applications: 1) an Initial QC, which relies exclusively on the first two non-invasive criteria (A and B) to determine eligibility of the organoids before entering a study (pre-study QC), and 2) a Final QC based on all the scoring criteria for a complete analysis at the end of a study (post-study QC) (Fig. 1A). Minimal thresholds have also been determined for passing the Initial and Final QC (Fig. 1B).

To validate this QC scoring methodology, we subjected cortical organoids at 60 days of culture to increasing doses of hydrogen peroxide (H_2_O_2_), a chemical known to induce apoptosis and oxidative stress, to generate organoids with varying quality levels (Fig. 1C). In this context, organoids were first selected for the H_2_O_2_ treatment experiment within a batch of cortical organoids, using the Initial QC method. H_2_O_2_ exposures were followed by a recovery period of one week, after which the exposed and non-exposed cortical organoids were evaluated for post-treatment quality using the complete Final QC.

### Initial Quality Control Scoring Streamlines the Selection of Cortical Organoids Based On Non-Invasive Criteria

By day 60 of culture, organoids exhibited spontaneous variability in quality due to the intrinsic differentiation heterogeneity and stochasticity within organoids (Fig. 2A). Consequently, we evaluated cortical organoids through the Initial QC based on morphology and size evolutions, to select those eligible for the H_2_O_2_ exposure experiment. Regarding the morphology evaluation (criterion A), the first index A1 referring to organoid density and compactness consistently achieved maximum scores of 5/5 across all the organoids (Fig. 2A, 2D). On the contrary, discrepancies were observed between the organoids for the second index A2 related to border integrity, with organoid #47 achieving the highest score of 5/5 (Fig. 2Ae, Fig. 2D), organoids #7 and #44 obtaining a score of 4/5 due to the presence of an area with less-defined border (Fig. 2Aa,d, Fig. 2D), and organoid #29 reaching a low score of 2/5 because of poorly-defined borders, but sufficient to pass the QC index (Fig. 2Ab, Fig.2D). However, organoid #31 failed to reach the minimum required score for border integrity (Fig. 2Ac, Fig. 2D), and was excluded from further analysis. Additionally, no cyst formation was observed, allowing all organoids to pass this third index A3 (Fig. 1A, D).

**Fig. 2:**
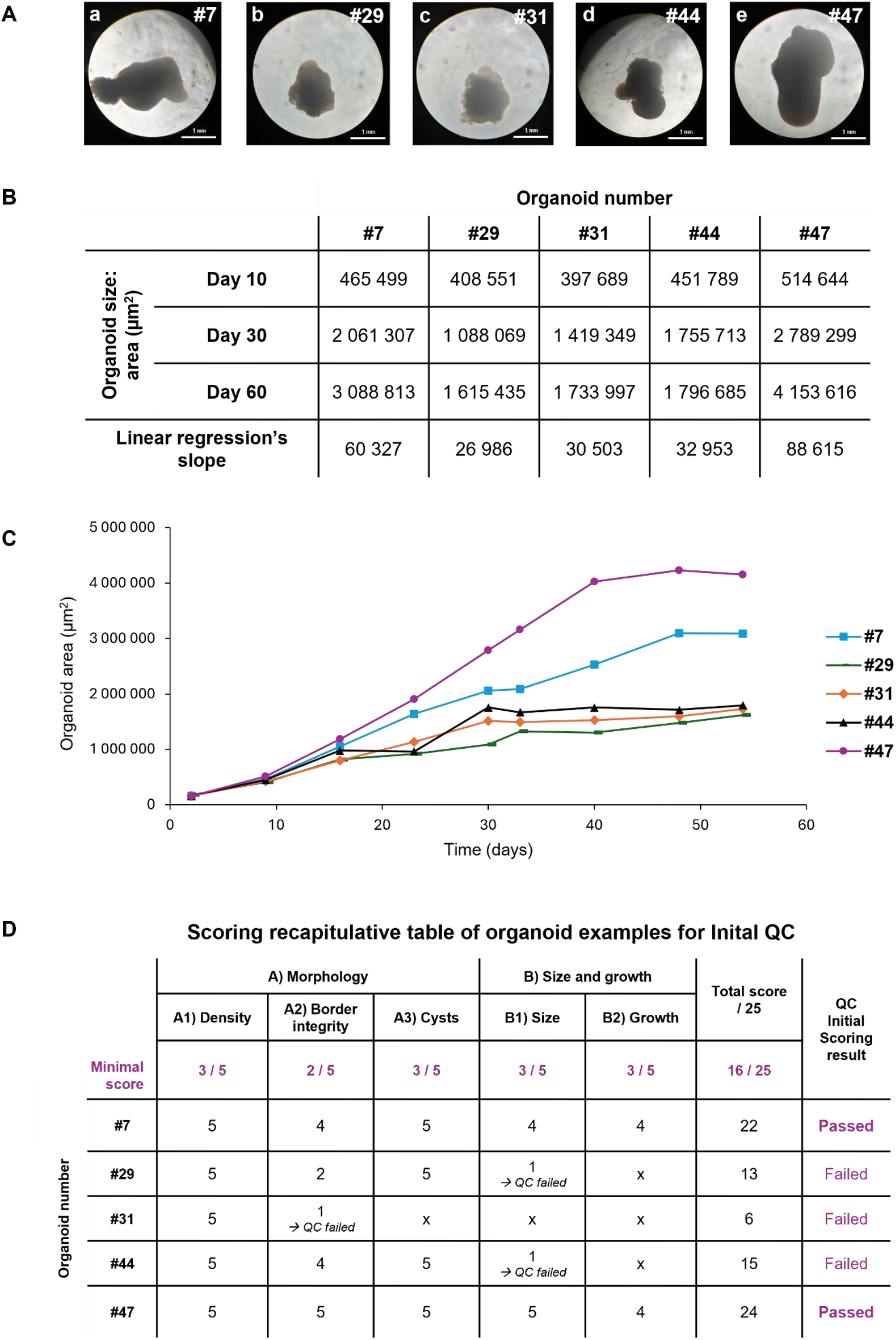
Quality Control (QC) for selection of cortical organoids before the H_2_O_2_ exposure experiment, following the Initial Scoring based on the first two non-invasive criteria Morphology and Growth evolutions. (A) Morphology of example cortical organoids within a batch at 60 days of culture (brightfield, 5X). (B) Table summarizing organoid sizes as surface areas at three timepoints of interest (Day 10, Day 30, and Day 60), as well as corresponding linear regression’s slopes. (C) Growth curves of individual organoids from D+2 to D+60 of culture. (D) Recapitulative table of scores obtained by the example organoids for each readout and indices of the Initial QC. Minimal scores per indices and the total minimal score required for Initial QC validation are mentioned. Results of QC for each organoid are indicated as Passed/Failed

Regarding the organoid sizes and growth evolutions (criterion B), organoids #29 and #44 were excluded due to insufficient growth both at day 60 and throughout the culture period (Fig. 2B, C, D). Interestingly, organoid #31 would not have passed this QC step as well but had already been excluded for the first criterion, emphasizing the relevance of this hierarchical order for QC evaluations. Consequently, only organoids #7 and #47 satisfied the minimum thresholds for the two non-invasive criteria and successfully passed the Initial QC (Fig. 2D). Overall, out of 58 cortical organoids generated within this batch, 10 were excluded due to a score lower than 16/25, representing 17% of the total population.

### Final Quality Control Scoring Effectively Evaluates Cortical Organoids Of Varying Quality Levels

Pre-selected cortical organoids via the Initial QC were included in the H_2_O_2_ exposure experiment to generate varying degrees of damage (Fig. 3, S2, S3). A total of six H_2_O_2_ concentrations were assayed (n = 4 organoids per group): 0% (untreated controls), 0.1%, 0.25%, 0.5%, 1%, and 5% H_2_O_2_. Before H_2_O_2_ exposures, all the organoids exhibited an optimal morphology, resulting from the Initial QC selection (Fig. 3a1-f1, S2a1-i1, S3a1-i1). After exposures, organoids exposed to 5% of H_2_O_2_ displayed severe loss of integrity and cellular disaggregation (Fig. 3f2, S3g2-i2), thus failing to pass the QC at the morphological criterion for the border integrity index (Tables 1, S2). This condition also prevented subsequent analyses based on organoid embedding and sectioning for immunostaining, therefore hampering further QC evaluation for these 5% H_2_O_2_-exposed organoids. Organoids treated with the other H_2_O_2_ concentrations succeeded to pass the morphology QC according to our criteria.

**Table 1:**
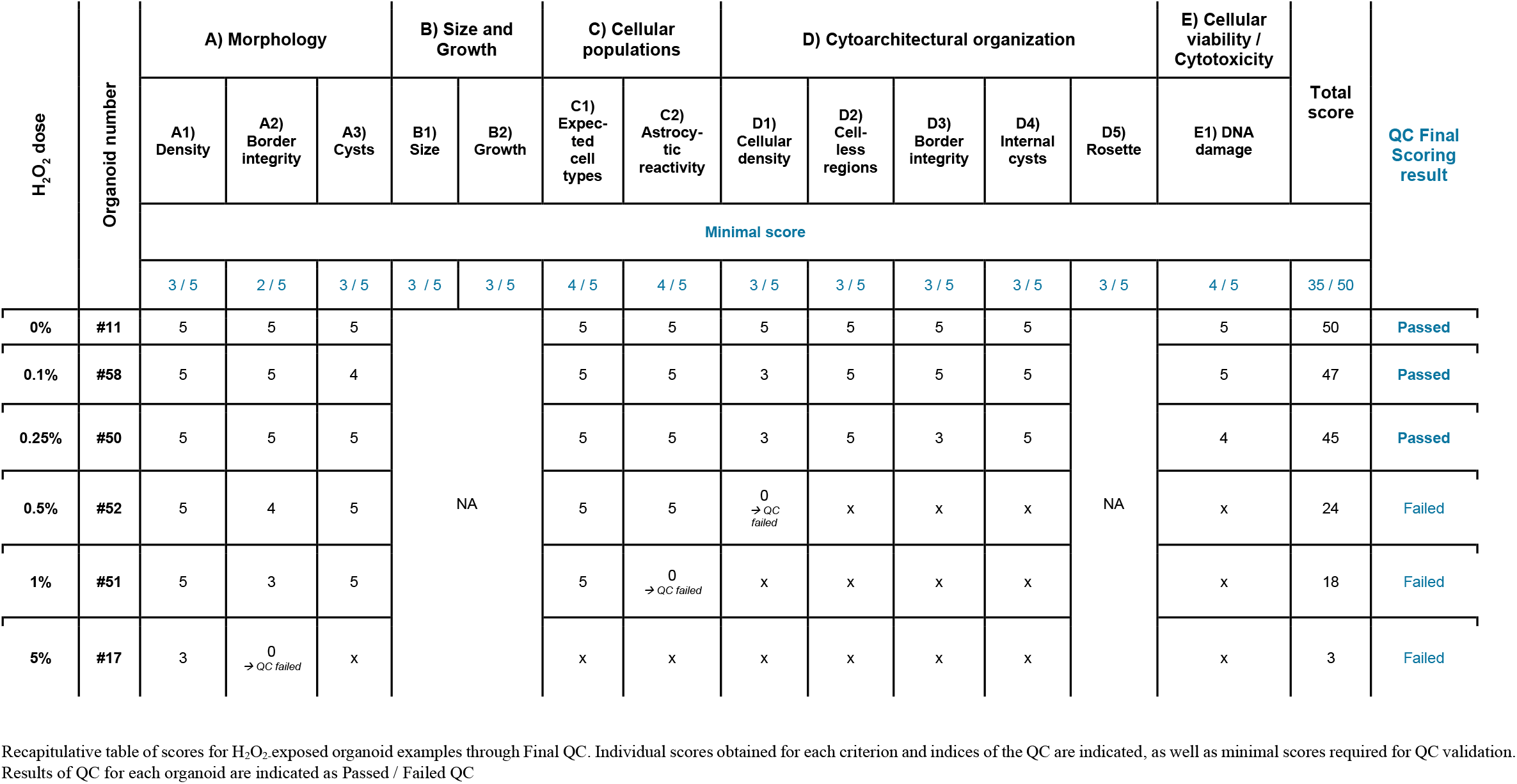
Scoring recapitulative table of H_2_O_2_-exposed organoid examples for Final QC.

**Fig. 3:**
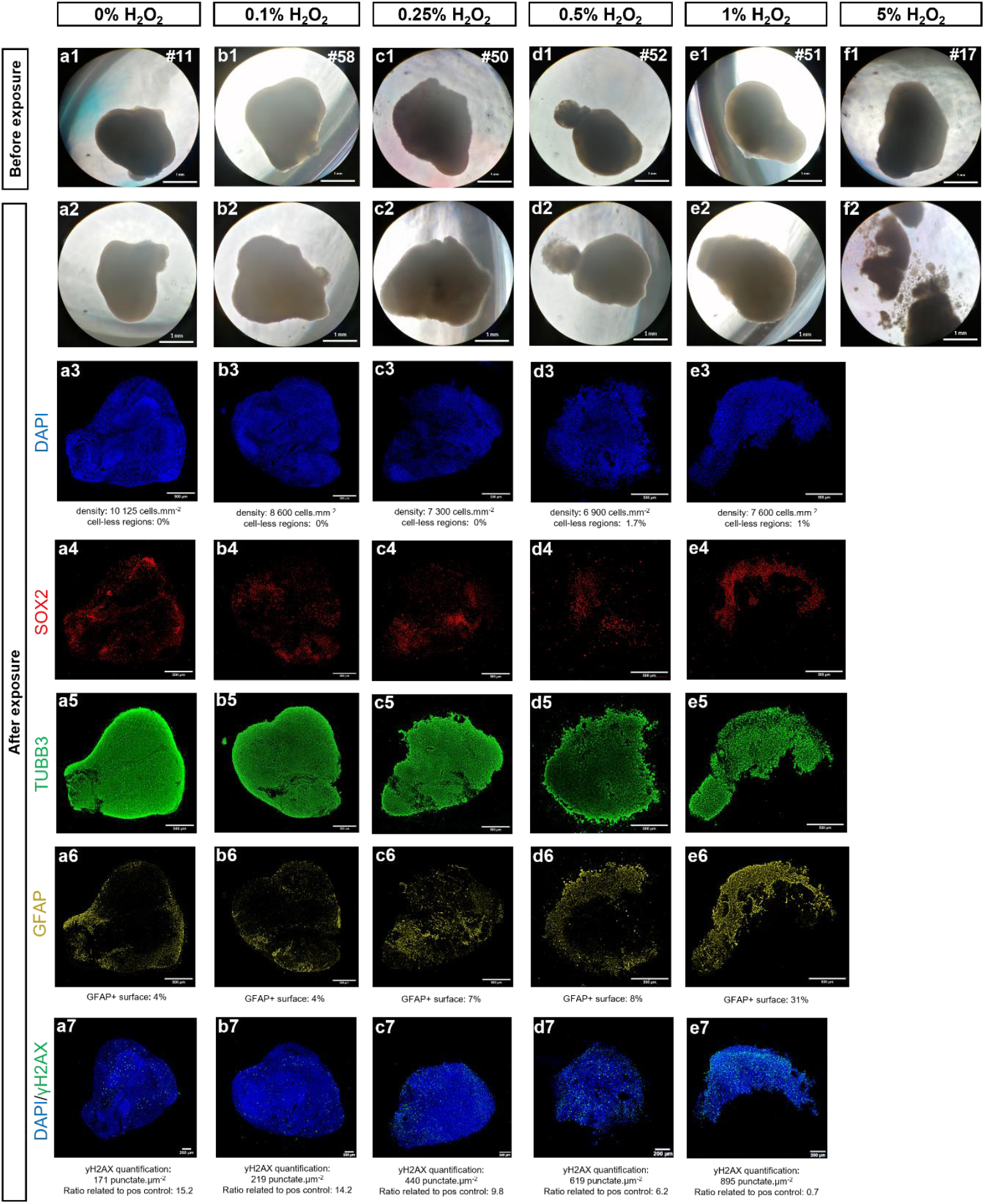
Quality Control (QC) for evaluation of cortical organoids after H_2_O_2_ exposures, following the Final Scoring based on all the criteria. H_2_O_2_ exposures on cortical organoids serve as examples of varying organoid quality levels through incremental H_2_O_2_ doses, with examples of organoids passing or not the QC. (a1-f1) Examples of cortical organoids exposed to different H_2_O_2_ doses comprised between 0% and 5%. Morphology before (a1-f1) and after (a2-f2) H_2_O_2_ exposures serve to evaluate the first criterion related to morphological quality evaluation (brightfield, 5X). Immunofluorescent staining of the example organoids for DAPI (a3-e3), neural progenitor marker SOX2 (a4-e4), neuronal marker TUBB3 (a5-e5), and astrocytic marker GFAP (a6-e6) enable the assessment of the following criteria: verification of cell types presence, assessment of Astrocytic reactivity, and evaluation of Cytoarchitectural organization. Immunofluorescent labeling of DNA damage with γH2AX marker enables evaluation of Cytotoxicity level (a7-e7) (Leica THUNDER microscope, 20X)

The size and growth profile criterion were not reassessed post-H_2_O_2_ exposures (Tables 1, S2), as the seven-day recovery period after H_2_O_2_ treatment was insufficient for meaningful growth analysis.

Subsequent invasive analyses were performed to evaluate cellular composition and cytoarchitectural organization within the organoids. Immunofluorescence staining confirmed the presence of neural progenitors (SOX2), immature neurons (TUBB3), and astrocytes (GFAP) across all remaining conditions (Fig. 3a3-e6, S2a3-i6, S3a3-d6), thus validating the QC criterion of cell type presence verification (Tables 1, S2). However, it must be noted that three organoids (#23, #30 and #47) could not be analyzed by immunolabeling, as they could not be embedded and sectioned with cryostat, likely due to a lack of compactness. Consequently, these organoids were excluded at this QC step (Table S2). Interestingly, they belonged to conditions where all the other organoids failed to pass the QC indices related to cellular composition and cytoarchitectural assessment (Tables 1, S2).

Regarding the astrocytic reactivity index, GFAP staining in the remaining organoids treated with 1% H_2_O_2_ (#51 and #53) suggested an excessive astrocytic reactivity, by covering 31% and 22% of the section area, respectively (Fig. 3e6, S3d6). This implies potential physiological disruption, thus excluding these organoids according to our QC criteria.

The next index evaluates the overall cellular density, based on DAPI staining analysis. Among the remaining organoids that have passed previous QC steps, we can observe that the cellular density was notably reduced in organoids exposed with 0.5% H_2_O_2_ (#49 and #52), with densities of 6 900 and 7 600 cells.mm^-2^, respectively (Fig. 3d3, S3b3), therefore not reaching the minimal threshold fixed in the scoring (Fig. S1) and leading to the exclusion of these organoids.

In contrast, no significant cytoarchitectural disruptions – such as the presence of large cell-less regions, severely altered borders, or occurrence of internal cysts – were observed in the remaining organoids, which therefore met the QC minimal standards for these criteria (Tables 1, S2). The rosette index was not assessed in this batch, as neurogenic niches were absent in the control organoids (Tables 1, S2).

Finally, cytotoxicity was evaluated via γH2AX staining, a marker of DNA double-stranded breaks, to quantify DNA damage. For all the remaining organoids exposed to 0%, 0.1% and 0.25% H_2_O_2_, the γH2AX quantification was significantly different from a positive control of maximal γH2AX labeling (Fig. 3a7-c7, S2a7-i7), therefore passing this final QC step (Tables 1, S2). Interestingly, all the organoids exposed to higher doses than 0.5% H_2_O_2_, that have been previously excluded at different QC steps, would have failed to pass this final criterion (Fig. 3d7,e7, S3a7-d7), confirming the validity and relevance of this hierarchical QC system.

Overall, Table 2 summarizes the individual final QC scoring results obtained by all the controls and exposed organoids, as well as the median scores reached per condition. We can observe that the unexposed controls and organoids exposed to the doses of H_2_O_2_ at 0.1% and 0.25% successfully passed the QC. However, organoids treated with the doses of 0.5%, 1% and 5% H_2_O_2_ failed to pass the QC evaluation. We can notice that these excluded organoids failed at different QC steps, following the hierarchical order of criteria, with 5% H_2_O_2_-exposed organoids excluded during the morphological criterion, 1% H_2_O_2_-exposed organoids failing regarding the cellular composition criterion, and 0.5% H_2_O_2_-exposed organoids rejected through either the cytoarchitectural or the cytotoxicity criteria. Similarly, median scores obtained per condition increase incrementally depending on the exposure doses, from a low score of 5/50 obtained for the highest dose of H_2_O_2_, up to an elevated score of 47/50 reached for the unexposed controls, thus correlating with the expected damage levels induced by the H_2_O_2_ graded exposures. These results demonstrate the sensitivity of this QC scoring methodology in efficiently and robustly distinguishing organoid quality levels.

**Table 2:**
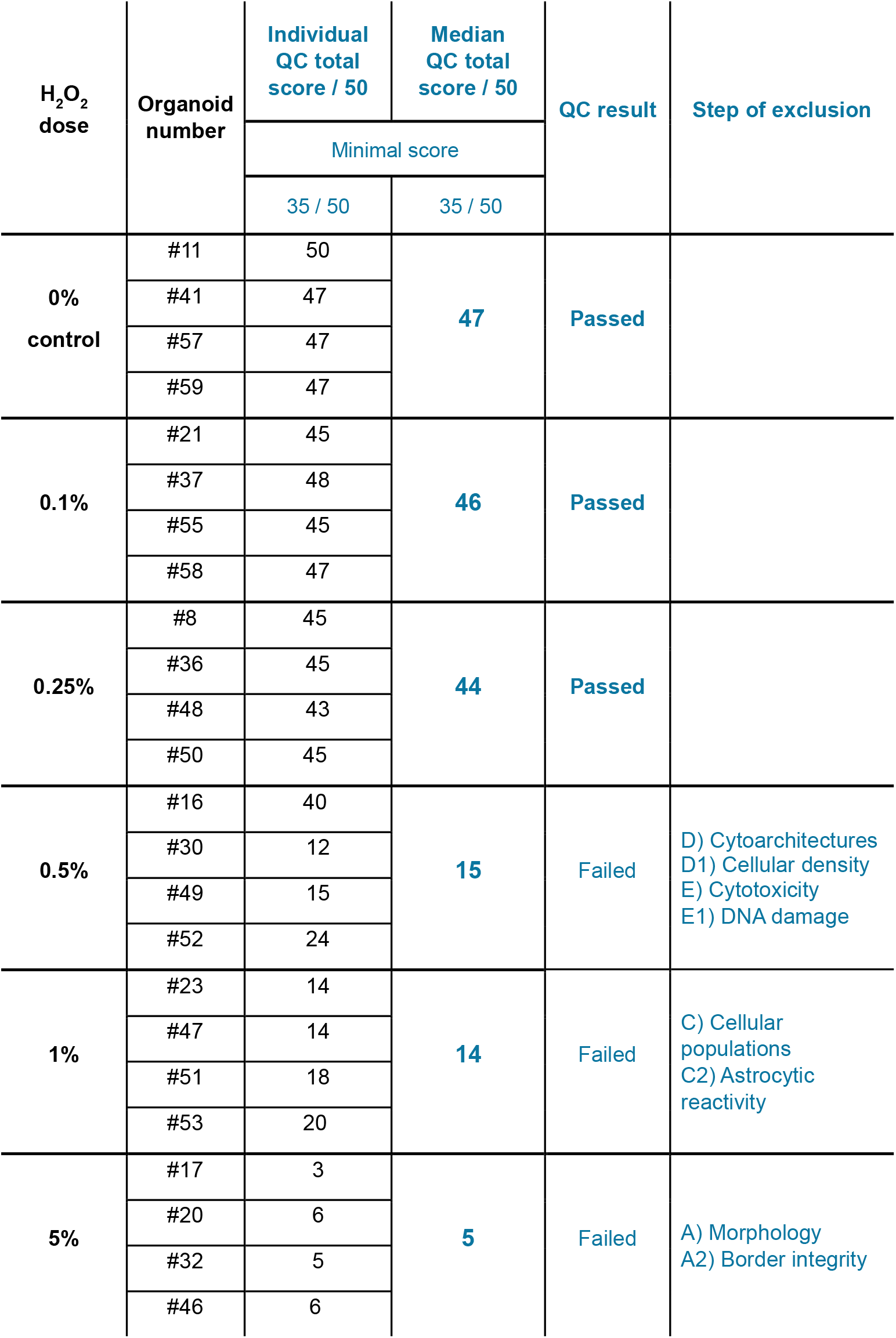
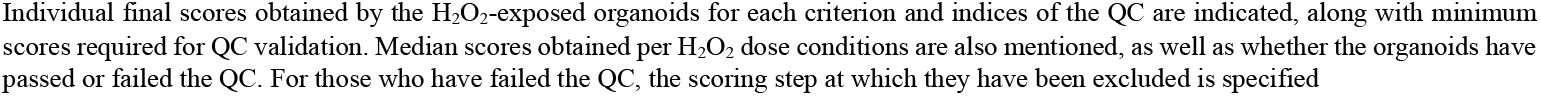
Recapitulative table of final QC scores obtained for H_2_O_2_-exposed cortical organoids.

| H<sub>2</sub>O<sub>2</sub><br>dose | Organoid<br>number | Individual<br>QC total<br>score / 50 | Median<br>QC total<br>score / 50 | QC result | Step of exclusion |
| --- | --- | --- | --- | --- | --- |
|  |  | Minimal score |  |  |  |
|  |  | 35 / 50 | 35 / 50 |  |  |
| 0%<br>control | #11 | 50 | 47 | Passed |  |
|  | #41 | 47 |  |  |  |
|  | #57 | 47 |  |  |  |
|  | #59 | 47 |  |  |  |
| 0.1% | #21 | 45 | 46 | Passed |  |
|  | #37 | 48 |  |  |  |
|  | #55 | 45 |  |  |  |
|  | #58 | 47 |  |  |  |
| 0.25% | #8 | 45 | 44 | Passed |  |
|  | #36 | 45 |  |  |  |
|  | #48 | 43 |  |  |  |
|  | #50 | 45 |  |  |  |
| 0.5% | #16 | 40 | 15 | Failed | D) Cytoarchitectures<br>D1) Cellular density<br>E) Cytotoxicity<br>E1) DNA damage |
|  | #30 | 12 |  |  |  |
|  | #49 | 15 |  |  |  |
|  | #52 | 24 |  |  |  |
| 1% | #23 | 14 | 14 | Failed | C) Cellular<br>populations<br>C2) Astrocytic<br>reactivity |
|  | #47 | 14 |  |  |  |
|  | #51 | 18 |  |  |  |
|  | #53 | 20 |  |  |  |
| 5% | #17 | 3 | 5 | Failed | A) Morphology<br>A2) Border integrity |
|  | #20 | 6 |  |  |  |
|  | #32 | 5 |  |  |  |
|  | #46 | 6 |  |  |  |

Individual final scores obtained by the H<sub>2</sub>O<sub>2</sub>-exposed organoids for each criterion and indices of the QC are indicated, along with minimum scores required for QC validation. Median scores obtained per H<sub>2</sub>O<sub>2</sub> dose conditions are also mentioned, as well as whether the organoids have passed or failed the QC. For those who have failed the QC, the scoring step at which they have been excluded is specified

## DISCUSSION

The stochastic nature of the differentiation of stem cells and their spontaneous self-organization within cerebral organoids lead to unpredictable variability among them [1]. Novel approaches have recently emerged to improve culture conditions and enhance organoid reproducibility. For instance, Brain Organoid-on-Chip systems, which rely on the use of microfluidic devices, offer precise fluid flow control and a more physiologically relevant microenvironment [45], improving cellular viability [35, 36, 61, 62], neural maturation [61, 63], and organoid homogeneity [61]. Other advancements include the use of bioreactors, that enable dynamic flow conditions and enhance oxygenation in large 3D cultures [64–66], as well as bioengineering strategies such as the use of 3D scaffolds and bioprinting [67–70], which further improve structural consistency. A critical gap remains despite these innovations: the absence of consensus on what defines high-quality cerebral organoids. This lack of standardized metrics or guidelines not only hinders meaningful comparisons across studies but also limits the broader applicability of these models. While complete uniformity should not be the goal, since no living systems are identical, excessive variability in cerebral organoid quality undermines their predictability and reproducibility, potentially leading to inaccurate or unreproducible findings, as well as wasted resources. This challenge is further exacerbated by the absence of standardized protocols for organoid generation, culture and characterization, with numerous in-lab adaptations. As highlighted in a recent publication on a consensus about cerebral organoid nomenclature [71], there is a pressing need for the establishment of standardized frameworks in the field. Altogether, this highlights the urgent necessity for a robust and user-friendly quality control framework to ensure cerebral organoid reliability in both academic and industrial applications.

Our scoring-based QC approach, adapted to 60-day cortical organoids, opens the way for a standardized quality control methodology. By incorporating multiple analysis criteria, including both qualitative macro- and micro-level observations, this framework provides a complete evaluation to lay the foundation for defining what constitutes a high-quality cortical organoid. Importantly, our proposed QC is structured hierarchically to rapidly exclude low-quality organoids, while reserving more detailed analyses for those that passed the initial parameters. This scoring system enables precise evaluation of each index and criterion using tailored examples and scoring scales, covering the full quality spectrum observed in cerebral organoids. While morphological criteria remain qualitative, the clarity and preciseness of the provided examples ensure robust evaluations. These illustrative examples enhance accessibility, allowing both experts and non-specialists to apply the scoring method effectively. For the other criteria, quantitative thresholds have been defined. Minimum scores have been established for each criterion, and failure to meet these scores immediately classifies the organoid as low quality, excluding it from further evaluations. If all the minimum scores are reached, additional thresholds incorporating multiple indices are applied to ensure a thorough quality assessment. As an example, for the morphology criterion, minimal scores required for the three indices are: 3/5 for density, 2/5 for border integrity, and 3/5 for absence of cysts, leading to a total of 8/15. However, the required minimal total score to pass the morphology criterion is 9/15, implying that the evaluated organoid could not obtain minimal quality levels for the three indices, and should reach a better quality for at least one of them (Fig. S1, 1B).

We demonstrated the effectiveness and robustness of our QC method through graded H_2_O_2_ exposures. Organoids were initially selected within a batch of cortical organoids for the exposure experiment using the Initial QC method. After H_2_O_2_ exposures, untreated and treated organoids were analyzed using the complete Final QC to assess post-treatment quality. Overall, the QC results demonstrated that only the untreated organoids exposed to 0.1% and 0.25% H_2_O_2_ passed the QC, while those exposed to higher doses above 0.5% did not (Table 2). Median scores reflected the severity of H_2_O_2_ exposure, ranging from a very low QC score (5/50) for the highest dose to an elevated score (47/50) for controls, correlating with the degree of damage caused by the graded H_2_O_2_ exposures. Importantly, failures occurred at different steps of the hierarchical QC process: organoids exposed to 5%, 1% and 0.5% H_2_O_2_ were excluded during the first criterion (morphology), the third criterion (cellular composition), or the fourth criterion (cytoarchitectures), respectively. These results emphasize the necessity of a stepwise evaluation, as certain defects are detectable only through deeper cellular or subcellular analysis. Taken together, these observations demonstrate the precision and reliability of the QC scoring system in differentiating organoid quality levels.

Interestingly, a few recent publications have demonstrated a growing interest in the field of cerebral organoids for the use of morphological criteria as reliable non-invasive readouts for the characterization of cerebral organoids [56–58]. Charles and colleagues have implemented a non-invasive quality control system relying on morphological criteria, enabling the classification of evaluated organoids in high- or low-quality categories for organoid pre-selection [58]. Remarkably, they integrated brightfield image processing with machine learning tools, opening the way for automated quality assessment. However, this system is based solely on morphological observations and does not account for other important analysis criteria that may provide valuable insights beyond what is visible at the macro-scale. As our study demonstrates, using a scoring scale enables the detection of subtle variations that a binary classification might overlook. Additionally, the inclusion of parameters such as sphericity can be questioned, as a brain is inherently far from spherical in shape. Filan et al. have also developed a non-invasive imaging analysis method for cerebral organoid characterization, also based on brightfield images, and considering several morphological criteria [56]. Using a 3D quantitative phase imaging technique, they can assess parameters in a non-invasive manner, such as cellular content, cell morphologies, and rosettes. Similarly, Ikeda and colleagues have implemented a non-invasive morphological characterization of cerebral organoids combined with transcriptomic analyses [57]. Interestingly, some analysis criteria are similar to those we selected, such as verification of transparency level and analysis of cystic structures [57]. Overall, morphological analysis serves as a valuable initial approach for assessing the quality of cerebral organoids. However, its findings should be validated through more detailed analyses, which often require invasive techniques.

While our proposed scoring system lays a foundation for organoid QC, there are still opportunities for further refinement and invasive techniques replacement.

It is important to note that this scoring system can be applied either manually, as proposed in this study, or by automation using image processing and machine learning analysis tools, offering flexibility, increased objectivity, and speed in execution as well as a gain in throughput. Automated analysis could enable organoid images to be processed and partially scored by the software, reducing variability in evaluations that might arise from individual interpretation. In particular, the automation could be envisaged primarily for criteria A) Morphology, based on brightfield images, as demonstrated by Charles and colleagues [58]; and D) Cytoarchitectures, based on immunofluorescence images.

Additionally, incorporating other non-invasive criteria could significantly enhance the transferability of the scoring system for pre-clinical applications. These could include the detection of specific markers in the conditioned medium, such as lactate dehydrogenase activity measurement for cytotoxicity evaluation [55], apoptosis quantification, measurement of reactive oxygen species for oxidative stress analysis, and evaluation of metabolic activity. While numerous ready-to-use kits are available on the market for these analyses, particular attention must be paid to the normalization step as these kits are typically designed to be normalized by cell number, which cannot be easily determined in 3D cultures [55].

Regarding the last criterion, which addresses cellular viability and cytotoxicity assessments, we evaluated DNA damage through γH2AX immunolabelling. However, other methods could also replace or complete this example, such as apoptosis detection via cleaved-caspase3 immunolabelling [53], TUNEL assay [52], or transcriptomic analyses evaluating pro- and anti-apoptotic markers BAX and BCL2 [51]. Ultimately, our framework provides flexibility, enabling the inclusion or exclusion of parameters based on the specific characteristics of the study (e.g., neurogenic niches, which could not be assessed here). However, to ensure consistency, it is crucial to define and validate thresholds through preliminary testing, especially when working with specific cell lines or different maturation timing. Similarly, growth dynamics should be adjusted according to the number of cells used during 3D seeding.

This QC system holds significant potential for biomedical research, ranging from fundamental to pre-clinical studies. For neurotoxicity studies, it could facilitate systematic comparisons between exposed and non-exposed organoids, using consistent scoring across criteria. Similarly, for disease modeling, the scoring system could be adapted to focus on specific phenotypes critical for recapitulating pathological hallmarks. By addressing these evolving needs, this framework paves the way for more robust, reproducible, and versatile organoid-based research. Notably, it represents a critical step toward the much-needed collaborative effort to define and standardize quality expectations for different types of organoids. As the field moves toward increasingly complex models, such as assembloids [72], maintaining scientific rigor requires a shared foundation.

## ACKNOWLEDGMENTS

The authors would like to express their sincere gratitude to François Hémard for his contributions to the acquisition, processing, and analysis of γH2AX images, and to Axel Fontanier for his valuable help in brightfield image acquisition. Special thanks are extended to Thomas Lemonnier for his work on reprogramming the BJ fibroblast cell line into induced pluripotent stem cells and to Elise Delage for her expert guidance on image analysis and fluorescence quantification. Finaly, we warmly thank Florian Larramendy for his review of the manuscript.

## STATEMENTS & DECLARATIONS

### Funding

The authors declare that no funds, grants, or other support were received during the preparation of this manuscript.

### Competing Interests

Financial interests: Héloïse Castiglione, Camille Baquerre, Benoît Guy Christian Maisonneuve, Jessica Rontard and Thibault Honegger are employed by NETRI company, whose CEO is Thibault Honegger.

### Author Contributions

All authors contributed to the study conception and design. Material preparation, data collection and analysis were performed by Héloïse Castiglione, Lucie Madrange, Camille Baquerre, Jessica Rontard and Pierre-Antoine Vigneron. The first draft of the manuscript was written by Héloïse Castiglione, Jessica Rontard and Pierre-Antoine Vigneron and all authors commented on previous versions of the manuscript. All authors read and approved the final manuscript.

### Data Availability

The data underlying this article will be shared on reasonable request to the corresponding authors.

## Notes

### Competing Interest Statement

H Castiglione, C Baquerre, BGC Maisonneuve, J Rontard and T Honegger are employed by NETRI company, whose CEO is T Honegger.

