## Supplementary material for "A robust and comprehensive quality control of cerebral cortical organoids: methodology and validation": Supp-Fig-S1

### Quality Control of 60-day Cortical Organoids

**Scoring scale:** 5 4 3 2 1 0

→ **5: high quality** (most optimal characteristics)
→ **0: poor quality** (suboptimal characteristics)

#### A) Morphology

- **Analytical method:** brightfield microscopy

##### A1) Overall color and structural density/compactness

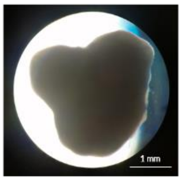

☑ uniform dark brown + high density

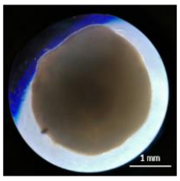

☑ uniform lighter brown + high density

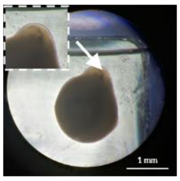

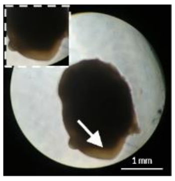

☑ dark brown, with light-brown borders (arrows) + high density

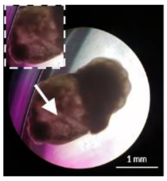

☑ dark/dense core + lighter/less dense protrusions (arrow)

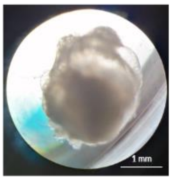

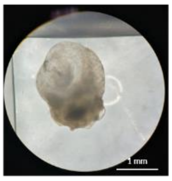

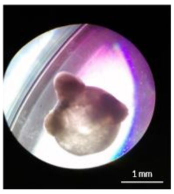

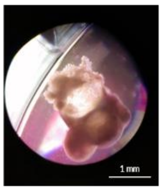

☒ lighter brown + low density/compactness (visible cell-less areas)

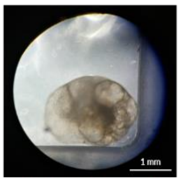

☒ light brown + very poor density

5 — 3 — 1 — 0

Minimal score to pass QC: ≥ 3

##### A2) Border integrity

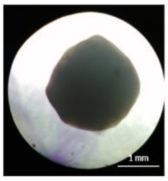

☑ well-defined borders, without cell shedding / detaching

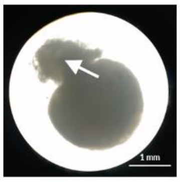

☑ defined borders, with some areas having +/- defined borders (arrow)

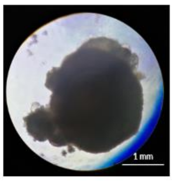

☑ +/- defined borders, with some areas having cell shedding

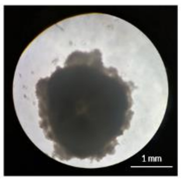

☑ not well-defined borders + cell shedding / detaching

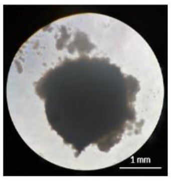

☒ poorly-defined borders + high cell detachment

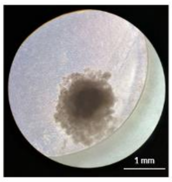

☒ very poorly-defined borders + high cell detachment + loss of integrity

5 — 4 — 3 — 2 — 1 — 0

Minimal score to pass QC: ≥ 2

##### A3) Presence/absence of cysts

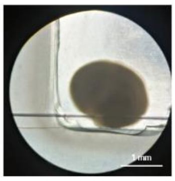

☑ absence of cysts

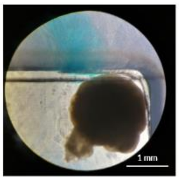

☑ presence of little + quite dense cyst(s)

☑ cyst area ≤ 10% of total organoid surface

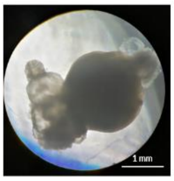

☑ presence of sparsely-dense cyst(s)

☑ 10% < cyst area ≤ 30% of total surface

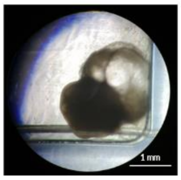

☒ presence of large + very poorly-dense cyst(s)

☒ 30% < cyst area ≤ 60% of total surface

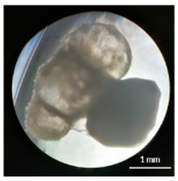

☒ presence of large + very poorly-dense cyst(s)

☒ 60% < cyst area ≤ 90% of total surface

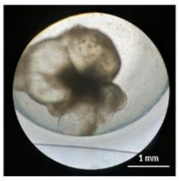

☒ presence of multiple cysts

☒ area of cysts > 90% of total surface

5 — 4 — 3 — 2 — 1 — 0

Minimal score to pass QC: ≥ 3

#### B) Size and growth evolution

- **Analytical method:** organoid surface area measurement, based on brightfield images

##### B1) Size

- Timeline of cortical organoid culture until D+60, with expected ranges of surface areas according to culture timepoints:

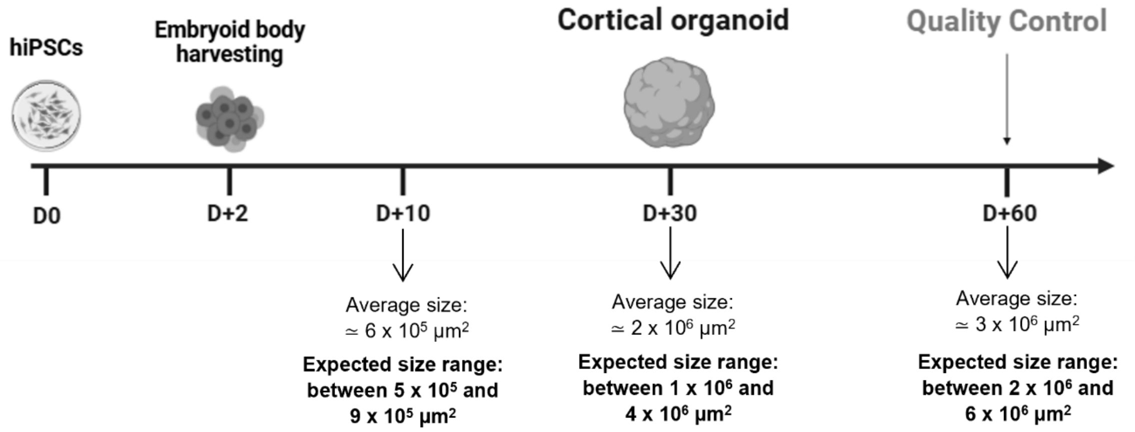


- ☑ size is comprised in all of the 3 ranges — **5**
- ☑ size is comprised in 2 ranges, including D+60 — **4**
- ☑ size is comprised in D+60 range only, not in the other ranges — **3**
- ☒ size is comprised in 2 ranges out of 3, not in D+60 range — **2**
- ☒ size is comprised in 1 range out of 3, not in D+60 range — **1**
- ☒ size is not comprised in any of the ranges — **0**

**Minimal score to pass QC: ≥ 3**

##### B2) Overall growth profile

- Based on organoid size evolution over time (growth curve)
- Slope of the growth curve's trendline from the beginning- (D+2) to the end-of-culture (D+60) timepoints has to be calculated

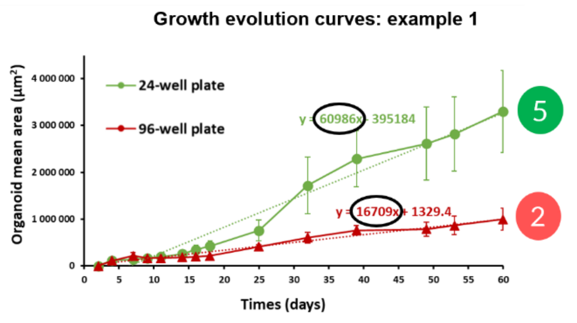


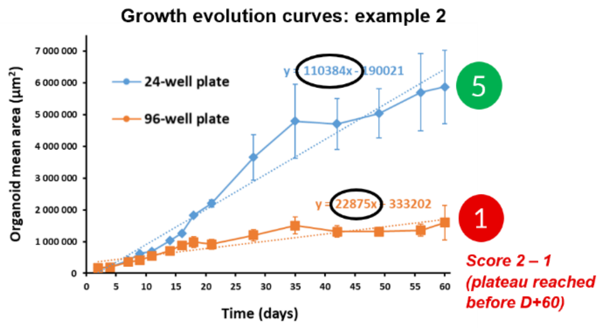


*24-well plate: cerebral organoids cultured in 24-well plate (500 µL – 1 mL of medium)*
*96-well plate: cerebral organoids cultured in 96-well plate (100 µL of medium)*

- ☑ slope ≥ 50 000 — **5**
- ☑ 40 000 ≤ slope < 50 000 — **4**
- ☑ 30 000 ≤ slope < 40 000 — **3**
- ☒ 20 000 ≤ slope < 30 000 — **2**
- ☒ 10 000 ≤ slope < 20 000 — **1**
- ☒ slope < 10 000 — **0**

☒ If a plateau is reached before D+60: deduct 1 point to final score

☒ If size decreases during growth: deduct 1 point to final score

**Minimal score to pass QC: ≥ 3**

#### C) Cellular populations

- **Analytical methods:** immunohistochemistry and / or transcriptomics

##### C1) Presence/absence of expected cell type markers at D+60

- **Neural progenitor markers:**

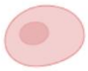

→ SOX2
→ PAX6
→ NESTIN

- **Neuronal markers:**

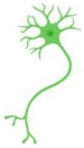

→ TUBB3
→ MAP2
→ NEUN
→ RELN

- **Astrocyte markers:**

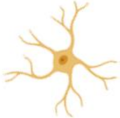

→ S100B
→ GFAP
→ SOX9

- ☑ presence of all 3 cell types — **5**
- ☑ presence of 2 cell types out of 3 (absence of neural progenitors) — **4**
- ☒ presence of 2 cell types out of 3 (absence of astrocytes) — **3**
- ☒ presence of 2 cell types out of 3 (absence of neurons) — **2**
- ☒ presence of 1 cell type out of 3 — **1**
- ☒ absence of any cell type — **0**

**Minimal score to pass QC: ≥ 4**

##### C2) Astrocytic reactivity

- **Based on Glial Fibrillary Acidic Protein (GFAP) positive surface expression related to total surface area**

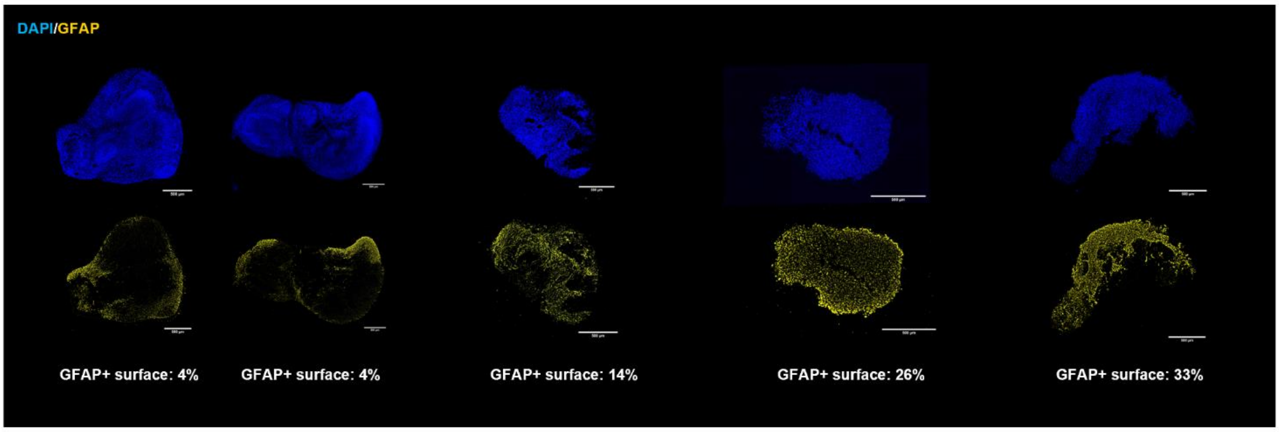


- ☑ GFAP+ surface < 10% of total surface — **5**
- ☑ GFAP+ surface between 10% and 20% of total surface — **4**
- ☒ GFAP+ surface between 20% and 30% of total surface — **1**
- ☒ GFAP+ surface > 30% of total surface — **0**

**Minimal score to pass QC: ≥ 4**

### D) Cytoarchitectural organization

- **Analytical method:** immunohistochemistry

#### D1) Cellular density

- **Based on DAPI positive surface related to total surface area (without considering cell-less areas/holes/necrotic cores)**
- **If less-densed areas are observed on organoid sections, the user should ensure that they are not technical artifacts resulting from the immunohistochemistry procedure**

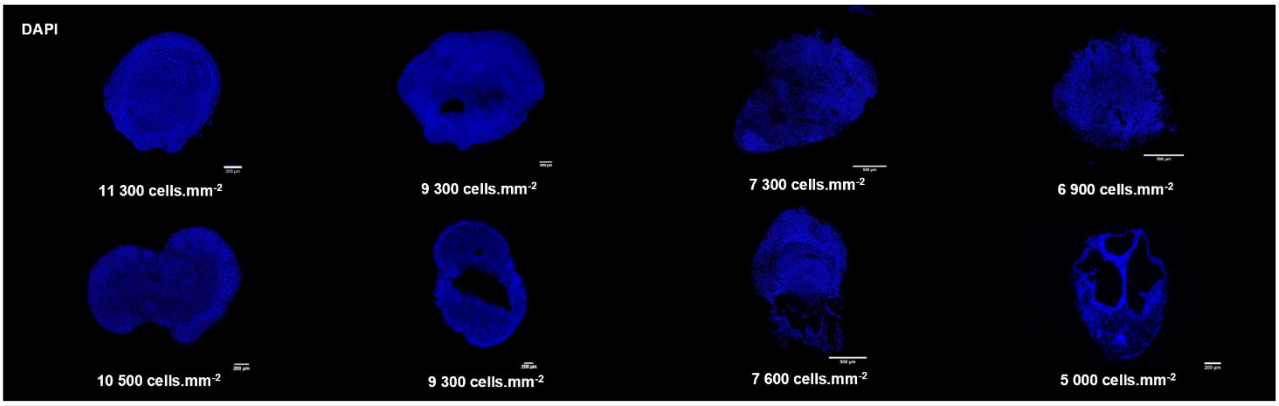


☑ high cellular density
☑ ≥ 10 000 cells.mm$^{-2}$

☑ quite high cellular density
☑ 9 000 ≤ between < 10 000 cells.mm$^{-2}$

☑ low cellular density (spaces between cells)
☑ 7 000 ≤ between < 9 000 cells.mm$^{-2}$

☒ very low cellular density
☒ < 7 000 cells.mm$^{-2}$

5 — 4 — 3 — 0

**Minimal score to pass QC: ≥ 3**

#### D2) Cell-less regions (e.g.: holes, necrotic cores):

- **Based on cell-less regions area measurement related to total surface area**
- **If cell-less regions are observed on organoid sections, the user should ensure that they are not technical artifacts resulting from the immunohistochemistry procedure**

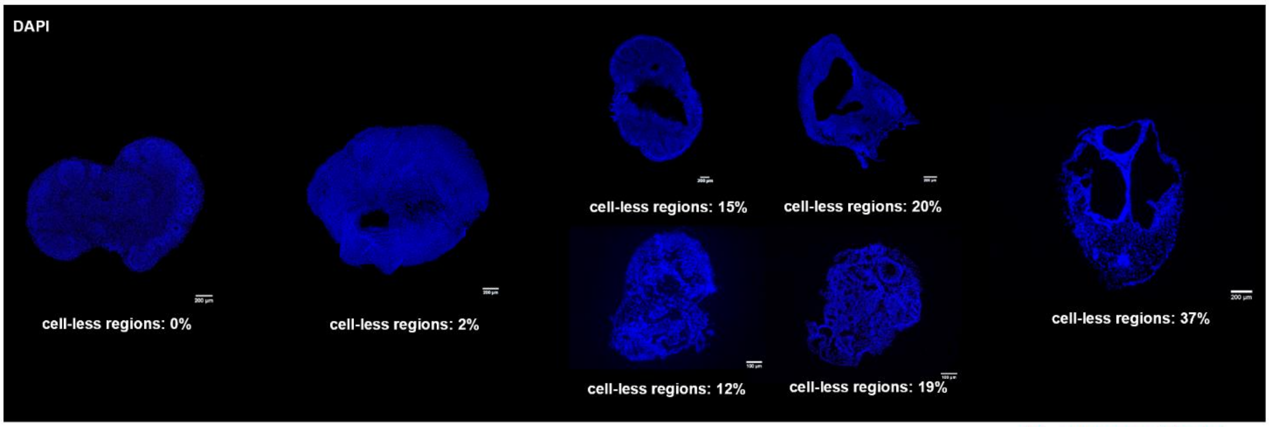


☑ 0% of cell-less regions

☑ presence of few / small holes
☑ < 5% of cell-less regions

☑ presence of larger holes
☑ 5% ≤ between < 20%

☒ presence of large / spread holes
☒ ≥ 20% of cell-less regions

5 — 4 — 3 — 0

**Minimal score to pass QC: ≥ 3**

#### D3) Border integrity

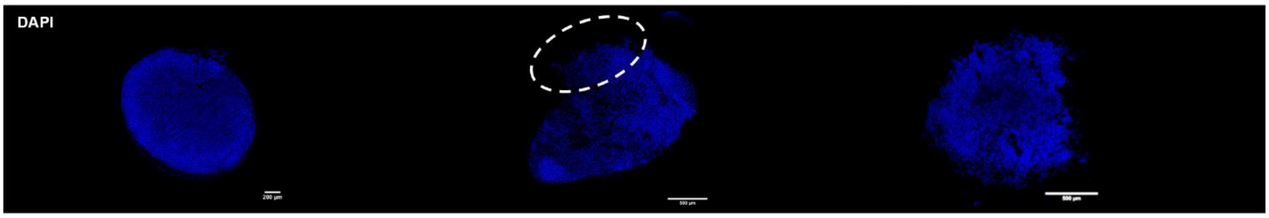


☑ well-defined border, without traces of cell detachment

☒ defined border, but some areas with cell detachment (circle)

☒ very poorly-defined border, with cell detachment

5 — 3 — 0

**Minimal score to pass QC: ≥ 3**

#### D) Cytoarchitectural organization

- **Analytical method:** immunohistochemistry

##### D4) Presence/absence of internal cysts

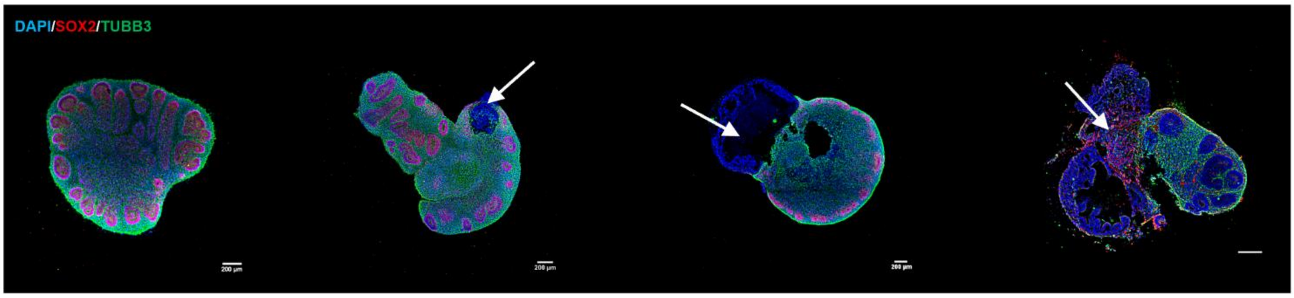


| 5 | 4 | 3 | 0 |
|---|---|---|---|
| ☑ absence of cysts | ☑ presence of small cyst (arrow)<br>☑ surface area ≤ 10% of total organoid surface | ☒ presence of large + very sparsely-dense cyst<br>☒ 10% < surface ≤ 30% of total surface | ☒ presence of very large and/or multiple cysts<br>☒ surface area > 30% |

**Minimal score to pass QC: ≥ 3**

##### D5) If applicable: Presence/absence & morphology/pattern of rosettes (neurogenic areas)

- Batch-dependent criterion: cortical organoid maturation timeline and speed vary between batches
- Criterion evaluated only if rosettes are observed in most of the cortical organoids within a batch

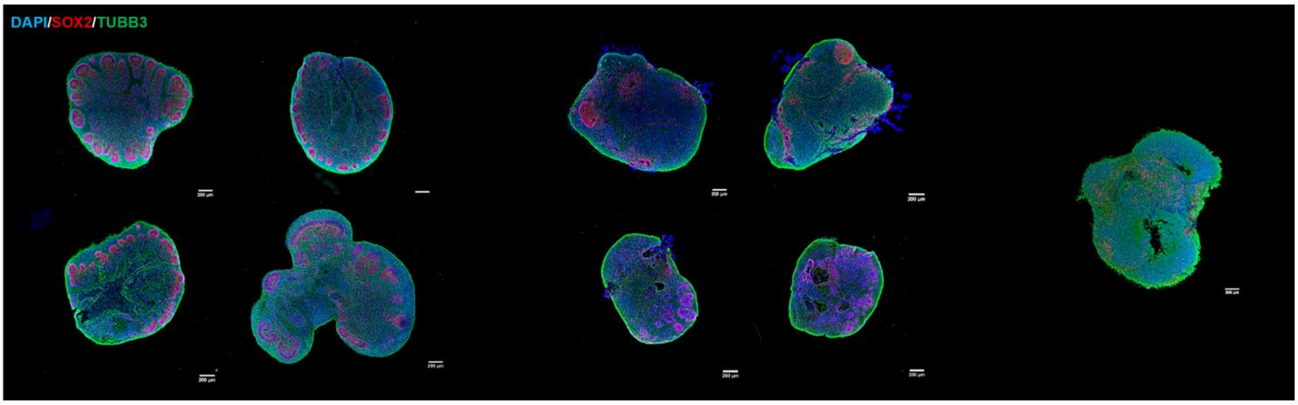


| 5 | 3 | 0 |
|---|---|---|
| ☑ presence of rosettes<br>☑ expected morphology (small and round-shaped)<br>☑ expected pattern at D+60 (on the organoid borders) | ☑ presence of rosettes<br>☑ +/- expected morphology and pattern (not well organized on the borders) | ☒ absence of rosettes |

**Minimal score to pass QC: ≥ 3**

#### E) Cellular viability/cytotoxicity evaluation

##### E1) Example: DNA damage evaluation based on γH2AX marker

- **Analytical method: immunohistochemistry**
- **γH2AX quantification: numeration of γH2AX immunolabeling, normalized to organoid surface area**
- **Results have to be compared with a γH2AX pos control: e.g. $H_2O_2$-exposed organoids (acute exposure with 1% $H_2O_2$ on 60-day cortical organoids, fixed at 7 days post-exposure)**
- **Statistical comparison based on the standard deviation of pos controls:**
    - Standard deviation calculation of pos controls ($\sigma_{control}$)
    - Difference calculation between the pos controls mean and the organoid value (Δ): $\Delta = \bar{x}_{control} - \text{organoid value}$
    - Ratio calculation between Δ and $\sigma_{control}$ : $\text{ratio} = \frac{\Delta}{\sigma_{control}}$
    - Comparison of ratio with predefined thresholds indicated in the scoring explanations:

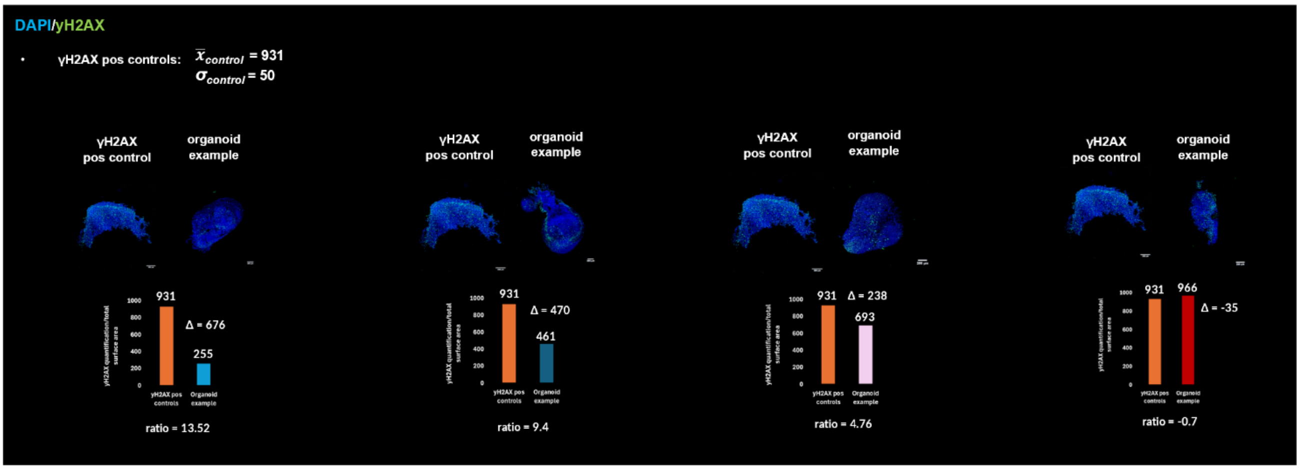


☑ γH2AX quantification lower than γH2AX pos controls

☑ with ratio ≥ 12 (i.e., a difference of at least 12 folds the standard deviation of γH2AX pos controls)

☑ γH2AX quantification lower than γH2AX pos controls

☑ with 9 ≤ ratio < 12

☒ γH2AX quantification lower than γH2AX pos controls

☒ but with ratio < 9

☒ γH2AX quantification higher than γH2AX pos controls

☒ (i.e., with negative ratio)

5 — 4 — 1 — 0

**Minimal score to pass QC: ≥ 4**

**Fig. S1:** Quality Control (QC) of 60-day cortical organoids, based on a scoring methodology. Detailed version of the QC scoring system, including the five primary criteria and sub-indices for organoid evaluation, accompanied by illustrative examples associated with score thresholds. Analytical methods and notes for certain criteria enable to facilitate the utilization and transposition of the scoring. The minimal required QC score for each index is also mentioned
