## Supplementary material for "A robust and comprehensive quality control of cerebral cortical organoids: methodology and validation": Supp-Fig-S2

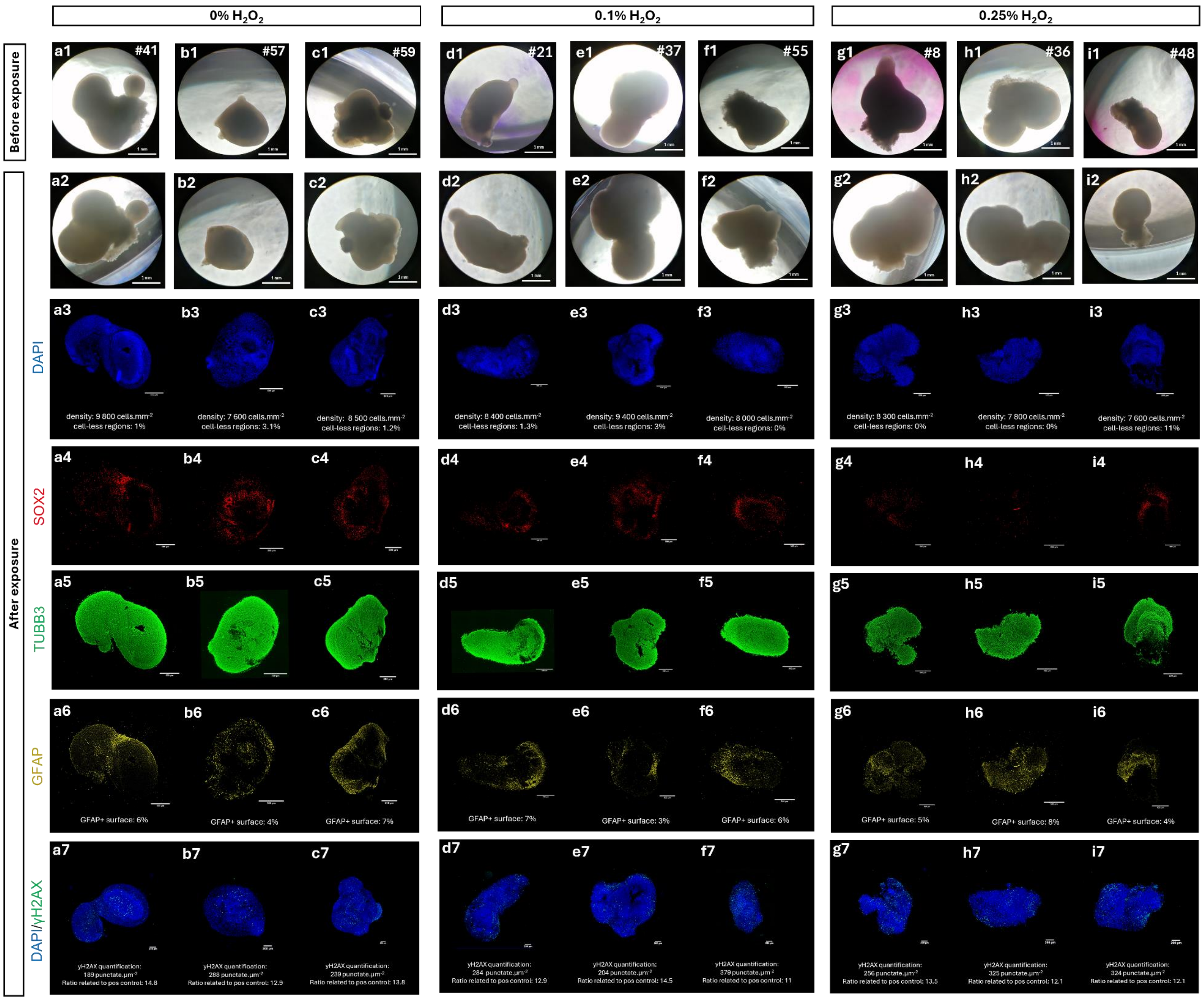
0% $H_2O_2$
0.1% $H_2O_2$
0.25% $H_2O_2$
Before exposure
After exposure
a1 #41
b1 #57
c1 #59
d1 #21
e1 #37
f1 #55
g1 #8
h1 #36
i1 #48
1 mm
a2
b2
c2
d2
e2
f2
g2
h2
i2
DAPI
a3
b3
c3
d3
e3
f3
g3
h3
i3
density: 9 800 cells.mm-2
cell-less regions: 1%
density: 7 600 cells.mm-2
cell-less regions: 3.1%
density: 8 500 cells.mm-2
cell-less regions: 1.2%
density: 8 400 cells.mm-2
cell-less regions: 1.3%
density: 9 400 cells.mm-2
cell-less regions: 3%
density: 8 000 cells.mm-2
cell-less regions: 0%
density: 8 300 cells.mm-2
cell-less regions: 0%
density: 7 800 cells.mm-2
cell-less regions: 0%
density: 7 600 cells.mm-2
cell-less regions: 11%
SOX2
a4
b4
c4
d4
e4
f4
g4
h4
i4
TUBB3
a5
b5
c5
d5
e5
f5
g5
h5
i5
GFAP
a6
b6
c6
d6
e6
f6
g6
h6
i6
GFAP+ surface: 6%
GFAP+ surface: 4%
GFAP+ surface: 7%
GFAP+ surface: 7%
GFAP+ surface: 3%
GFAP+ surface: 6%
GFAP+ surface: 5%
GFAP+ surface: 8%
GFAP+ surface: 4%
DAPI/γH2AX
a7
b7
c7
d7
e7
f7
g7
h7
i7
γH2AX quantification:
189 punctate.μm-2
Ratio related to pos control: 14.8
γH2AX quantification:
288 punctate.μm-2
Ratio related to pos control: 12.9
γH2AX quantification:
239 punctate.μm-2
Ratio related to pos control: 13.8
γH2AX quantification:
284 punctate.μm-2
Ratio related to pos control: 12.9
γH2AX quantification:
204 punctate.μm-2
Ratio related to pos control: 14.5
γH2AX quantification:
379 punctate.μm-2
Ratio related to pos control: 11
γH2AX quantification:
256 punctate.μm-2
Ratio related to pos control: 13.5
γH2AX quantification:
325 punctate.μm-2
Ratio related to pos control: 12.1
γH2AX quantification:
324 punctate.μm-2
Ratio related to pos control: 12.1

**Fig. S2:** Illustrative images of all cortical organoids after $H_2O_2$ exposures (first part of the figure, to be continued in Fig. S3). (a1-i1) Organoids exposed to different $H_2O_2$ doses comprised between 0% and 0.25%. Morphology before (a1-i1) and after (a2-i2) $H_2O_2$ exposures serve to evaluate the first criterion related to morphological quality evaluation (brightfield, 5X). Immunofluorescent staining for DAPI (a3-i3), neural progenitor marker SOX2 (a4-i4), neuronal marker TUBB3 (a5-i5), and astrocytic marker GFAP (a6-i6) enable the assessment of the following criteria: verification of Cell types presence, assessment of Astrocytic reactivity, and evaluation of Cytoarchitectural organization. Immunofluorescent labeling of DNA damage with yH2AX marker enables evaluation of Cytotoxicity level (a7-i7) (Leica THUNDER microscope, 20X). Additional data on the other $H_2O_2$ exposure doses can be found in Fig. S3
