## Supplementary material for "A robust and comprehensive quality control of cerebral cortical organoids: methodology and validation": Supp-Fig-S3

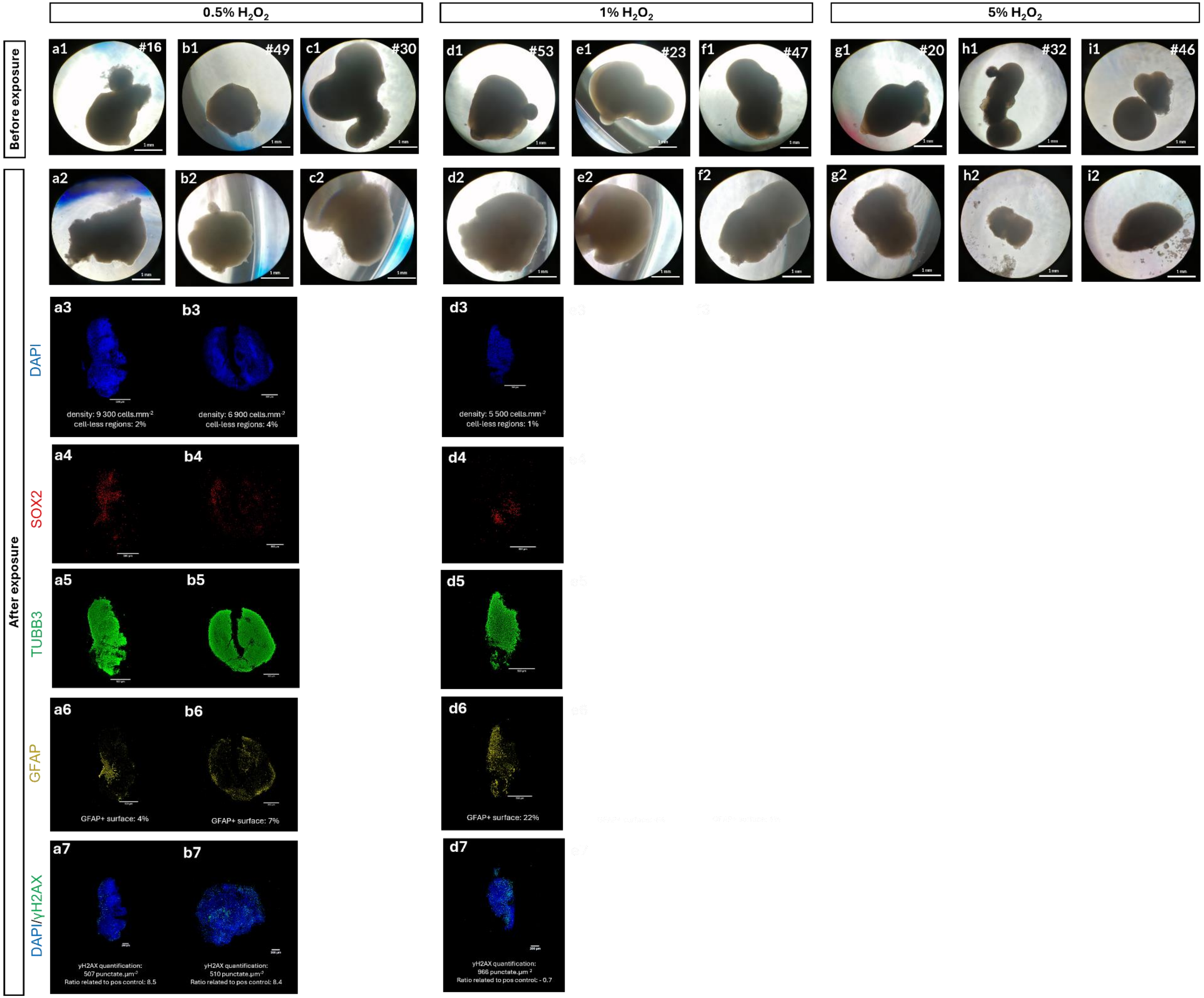
0.5% H2O2
1% H2O2
5% H2O2
Before exposure
After exposure
a1 #16
b1 #49
c1 #30
d1 #53
e1 #23
f1 #47
g1 #20
h1 #32
i1 #46
a2
b2
c2
d2
e2
f2
g2
h2
i2
DAPI
a3
b3
d3
density: 9 300 cells.mm-2
cell-less regions: 2%
density: 6 900 cells.mm-2
cell-less regions: 4%
density: 5 500 cells.mm-2
cell-less regions: 1%
SOX2
a4
b4
d4
TUBB3
a5
b5
d5
GFAP
a6
b6
d6
GFAP+ surface: 4%
GFAP+ surface: 7%
GFAP+ surface: 22%
DAPI/γH2AX
a7
b7
d7
γH2AX quantification:
507 punctate.μm-2
Ratio related to pos control: 8.5
γH2AX quantification:
510 punctate.μm-2
Ratio related to pos control: 8.4
γH2AX quantification:
966 punctate.μm-2
Ratio related to pos control: - 0.7

**Fig. S3:** Illustrative images of all cortical organoids after $H_2O_2$ exposures (second part of the figure). Quality Control (QC) for evaluation of cortical organoids after $H_2O_2$ exposures, following the Final Scoring based on all the criteria. (a1-i1) Organoids exposed to different $H_2O_2$ doses comprised between 0.5% and 5%. Morphology before (a1-i1) and after (a2-i2) $H_2O_2$ exposures serve to evaluate the first criterion related to morphological quality evaluation (brightfield, 5X). Immunofluorescent staining for DAPI (a3-d3), neural progenitor marker SOX2 (a4-d4), neuronal marker TUBB3 (a5-d5), and astrocytic marker GFAP (a6-d6) enable the assessment of the following criteria: verification of Cell types presence, assessment of Astrocytic reactivity, and evaluation of Cytoarchitectural organization. Immunofluorescent labeling of DNA damage with yH2AX marker enables evaluation of Cytotoxicity level (a7-d7) (Leica THUNDER microscope, 20X). Organoids for which the immunofluorescent staining images are not presented correspond to those that could not undergo embedding and cryosectioning processing due to insufficient density and compactness. Additional data on the other $H_2O_2$ exposure doses can be found in Fig. S2
