## Supplementary material for "A robust and comprehensive quality control of cerebral cortical organoids: methodology and validation": Supp-Fig-S4

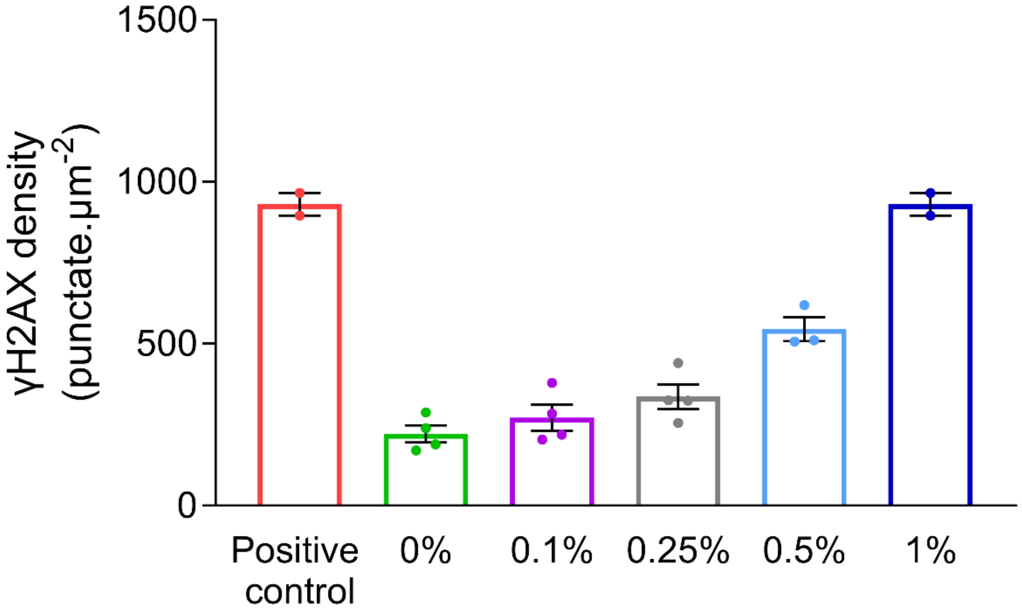


**Fig. S4:** Cytotoxicity evaluation index for $H_2O_2$-exposed organoids. Bar plot representing the γH2AX quantification in the organoids exposed to varying $H_2O_2$ concentrations between 0% and 5%. γH2AX quantification is expressed in punctate.µm2 (n=4 organoid per condition, ± SEM)
