## Supplementary material for "A robust and comprehensive quality control of cerebral cortical organoids: methodology and validation": Supp-Table-S1

**Supplementary Table S1: Primary and secondary antibodies used for immunofluorescent staining**

| Primary Antibodies | Host and Isotype | Reference and Supplier | Final dilution |
|---|---|---|---|
| Anti-SOX2 | Mouse IgG1 | 66411-1-Ig, Proteintech | 1:500 |
| Anti-TUBB3 | Mouse IgG2a | 801202, Biolegend | 1:500 |
| Anti-GFAP | Rabbit IgG | Z0334, Dako | 1:1000 |
| Anti-yH2AX | Mouse IgG1 | 05-636, Sigma-Aldrich | 1:500 |
| **Secondary Antibodies** | **Host** | **Reference and Supplier** | **Final dilution** |
| Alexa Fluor® Cy3 anti-Mouse IgG1 | Goat | 115-165-205, Jackson ImmunoResearch | 1:500 |
| Alexa Fluor® 488 anti-Mouse IgG2a | Goat | 115-547-186, Jackson ImmunoResearch | 1:500 |
| Alexa Fluor® 647 anti-Rabbit IgG | Goat | 111-605-144, Jackson ImmunoResearch | 1:500 |
| Alexa Fluor® 488 anti-Mouse IgG1 | Goat | 115-547-185, Jackson ImmunoResearch | 1:500 |
