## Supplementary material for "A robust and comprehensive quality control of cerebral cortical organoids: methodology and validation": Supp-Table-S2

### Supplementary Table S2: Recapitulative table of QC scores obtained for $H_2O_2$-exposed cortical organoids

| $H_2O_2$ dose | Organoid number | A) Morphology | | | B) Size and Growth | | D) Cellular populations | | D) Cytoarchitectural organization | | | | | E) Cellular viability / Cytotoxicity | Total score | QC Final Scoring result |
|---|---|---|---|---|---|---|---|---|---|---|---|---|---|---|---|---|
| | | A1) Density | A2) Border integrity | A3) Cysts | B1) Size | B2) Growth | C1) Expected cell types | C2) Astrocytic reactivity | D1) Cellular density | D2) Cell-less regions | D3) Border integrity | D4) Internal cysts | D5) Rosettes | E1) DNA damage | | |
| | | Minimal score | | | | | | | | | | | | | | |
|  |  | 3 / 5 | 2 / 5 | 3 / 5 | 3 / 5 | 3 / 5 | 4 / 5 | 4 / 5 | 3 / 5 | 3 / 5 | 3 / 5 | 3 / 5 | 3 / 5 | 4 / 5 | 35 / 50 |  |
| 0% | #11 | 5 | 5 | 5 | | | 5 | 5 | 5 | 5 | 5 | 5 | | 5 | 50 | **Passed** |
| | #41 | 5 | 4 | 5 | | | 5 | 5 | 4 | 4 | 5 | 5 | | 5 | 47 | **Passed** |
| | #57 | 5 | 5 | 5 | | | 5 | 5 | 3 | 4 | 5 | 5 | | 5 | 47 | **Passed** |
| | #59 | 5 | 5 | 5 | | | 5 | 5 | 3 | 4 | 5 | 5 | | 5 | 47 | **Passed** |
| 0.1% | #21 | 5 | 5 | 5 | | | 5 | 5 | 3 | 4 | 3 | 5 | | 5 | 45 | **Passed** |
| | #37 | 5 | 5 | 5 | | | 5 | 5 | 4 | 4 | 5 | 5 | | 5 | 48 | **Passed** |
| | #55 | 5 | 4 | 5 | | | 5 | 5 | 3 | 5 | 3 | 5 | | 5 | 45 | **Passed** |
| | #58 | 5 | 5 | 4 | | | 5 | 5 | 3 | 5 | 5 | 5 | | 5 | 47 | **Passed** |
| 0.25% | #8 | 5 | 4 | 5 | | | 5 | 5 | 3 | 5 | 3 | 5 | | 5 | 45 | **Passed** |
| | #36 | 5 | 4 | 5 | | | 5 | 5 | 3 | 5 | 3 | 5 | | 5 | 45 | **Passed** |
| | #48 | 5 | 4 | 5 | NA | | 5 | 5 | 3 | 3 | 3 | 5 | NA | 5 | 43 | **Passed** |
| | #50 | 5 | 5 | 5 | | | 5 | 5 | 3 | 5 | 3 | 5 | | 4 | 45 | **Passed** |
| 0.5% | #16 | 5 | 3 | 5 | | | 5 | 5 | 4 | 4 | 3 | 5 | | *1 → QC failed* | 40 | Failed |
| | #30 | 5 | 2 | 5 | | | *NA → QC failed* | x | x | x | x | x | | x | 12 | Failed |
| | #49 | 5 | 5 | 5 | | | 5 | 5 | *0 → QC failed* | x | x | x | | x | 15 | Failed |
| | #52 | 5 | 4 | 5 | | | 5 | 5 | *0 → QC failed* | x | x | x | | x | 24 | Failed |
| 1% | #23 | 5 | 4 | 5 | | | *NA → QC failed* | x | x | x | x | x | | x | 14 | Failed |
| | #47 | 5 | 4 | 5 | | | *NA → QC failed* | x | x | x | x | x | | x | 14 | Failed |
| | #51 | 5 | 3 | 5 | | | 5 | *0 → QC failed* | x | x | x | x | | x | 18 | Failed |

| | | | | | | | | | | | | | | | |
|---|---|---|---|---|---|---|---|---|---|---|---|---|---|---|---|
| | **#53** | 5 | 4 | 5 | | 5 | *1 → QC failed* | x | x | x | x | | x | 20 | Failed |
| **5%** | **#17** | 3 | *0 → QC failed* | x | | x | x | x | x | x | x | | x | 3 | Failed |
| | **#20** | 5 | *1 → QC failed* | x | | x | x | x | x | x | x | | x | 6 | Failed |
| | **#32** | 5 | *0 → QC failed* | x | | x | x | x | x | x | x | | X | 5 | Failed |
| | **#46** | 5 | *0 → QC failed* | x | | x | x | x | x | x | x | | X | 5 | Failed |

Summary table of scores obtained by all the $H_2O_2$-exposed cortical organoids for each criterion and indices of the QC. Minimal scores per indices and total minimal score required for QC validation are mentioned. Individual final QC scores obtained for each organoid, as well as median scores obtained per $H_2O_2$ dose conditions, are indicated. In addition, the final QC result is mentioned as QC passed/failed
